# Assembly of hierarchical multiphase condensates using designer surfactant proteins

**DOI:** 10.1101/2024.12.11.628011

**Authors:** Muyang Guan, Daniel A. Hammer, Matthew C. Good

## Abstract

Biomolecular condensates are complex fluids formed by liquid-liquid phase separation of macromolecules. Similar to other types of soft matter, they feature a range of biophysical properties that distinguish them from the cellular milieu. As a separate phase, they have an identifiable interface that dictates their interaction with the cytoplasm and other membraneless organelles. In this work, we engineer the interface of condensates to build novel hierarchical mesoscale structures from two immiscible disordered proteins: the RGG domain of LAF-1, a RNA-processing protein involved in germ granule assembly and the low complexity domain (LC) of FUS, an RNA binding protein whose aggregation is implicated in age-related neurodegeneration. RGG and FUS LC do not co-partition with one another and instead form discrete protein-rich condensed phases. Despite their apparent immiscibility, we identified conditions that can promote hierarchical assembly, either kinetically trapping one phase in the other, or using a designer surfactant protein that reduces the interfacial tension between the two phases. In addition, we studied factors that impact condensate miscibility and structure formation, including surface properties and viscoelasticity. This study probes the principles that underlie formation and assembly of complex structures from biomolecular condensates and provides a strategy for designing synthetic multiphasic materials capable of spatial partitioning.

## Introduction

Biomolecular condensates are multimolecular assemblies that form through liquid-liquid phase separation of macromolecules, including proteins and RNAs (1). Phase separation is mediated by various modes of homotypic interactions of protein amino acid side chains, such as charge-charge, cation-π, π-π interactions, and hydrogen bonding, as well as their interactions with nucleic acids (2-4). These condensates show typical rheological characteristics of complex fluids in vitro and in vivo, such as dripping, wetting, fusion, Ostwald ripening, surface tension, and viscoelasticity (1, 5-9). The rheological properties of condensates are linked to biological functions and aberrant phase transitions have disease implications (10-13).

Some cellular organelles such as the nucleolus are multiphasic condensates consisting of subcompartments that carry out distinct cellular functions (14). The multiphase structure is important for function, contributing to the transcription and vectorial processing of ribosomal RNAs. The components forming the membraneless organelle remain immiscible yet self-assemble into a functional hierarchical structure (14). The formation of such structures are governed by the interfacial energies of individual phases, such that the phases could be arranged in a manner that minimizes the system free energy (14).

Polymer model systems have been used to generate multiphase condensates. Self-assembly of oppositely charged polymers yields complex coacervates with multiphase architecture (15). For complex coacervates driven by charge interactions, critical salt concentrations govern polymer immiscibility. Further condensates can be generated from elastin-like polypeptides (ELPs), and diblock surfactants containing a hydrophilic and a hydrophobic block will stabilize the interface of a more hydrophobic condensate and prevent coarsening (16). Multiphase structures also observed for dynamically arrested condensates, which are capable or forming double-emusions and usually driven by an out-of-equilibrium process (17, 18).

A key feature of surfactants is their amphilicity which drives their localization to the interface between an aqeuous continuous phase and a condensed phase and that lowers surface tension (19). Kelley and coworkers synthesized a diblock protein surfactant that localizes at the interface of the condensate and continuous phase solvent and has functionality; it prevents droplets from fusing (19). However, the use of surfactants to modulate condensate mixing has yet to be tested for two immiscible protein condensed phases. RNAs can also function as surfactants (20). Interestingly, additional RNA can drive the immiscibility of two disordered peptides (21). To date, however, there has been little work done to tune interfacial tension between two peptide condensed phases and building hiearchical structures by engineering surfactant proteins.

Inspired by the existance of endogenous hiearchical organelles, we set out to build synthetic multiphase compartments from disordered proteins that form immiscible partially-wetting condensed phases. We previously identified an orthogonal pair of condensate-forming intrinsically disordered regions (IDRs) that show immiscibility in vitro and in cells (22). One IDR is the N-terminal 168 amino acid RGG region of C. elegans germ granule protein LAF-1, which is rich in arginines and glycines, and is necessary and sufficient for phase separation (5). Dimerization of RGG domain induces potent phase separation and can serve as the platform for the RGG based condensates capable of controlled assembly and dissolution, and the recruiting and release of cargo (23). The second IDR is FUS LC, the N-terminal 163 amino acids QGSY-rich domain of human FUS protein, which is also suffcient for phase separation (24) and is prone to aggregation and formation of amyloid fibrils involved in amyotrophic lateral sclerosis and frontotemporal dementia (10, 25). Unlike RGG, this domain lacks postively charged residues. RGG and FUS LC form discrete condensates – the scaffolds do not co-partition with one another – when reconstituted in vitro or when expressed in living cells, including yeast and mammalian cells (22). The selectivity of condensation is regulated by a balance between homotypic and heterotypic interactions. The native FUS LC does not mix with RGG; however, substituting 32 segregated charges (16 Arg + 16 Asp) into FUS LC induces heterotypic interactions with LAF-1 RGG, causing it to co-partition to RGG condensates (22). These results clearly highlight sequence-encoded immiscibility of the two disordered polypeptides which are capable of forming discrete condensed phases and offer a useful test case for our current experiments.

In this study, we sought to control hiearchical assembly of IDP phases built from FUS LC and LAF-1 RGG to generate novel mesoscale compartments. Such multiphase assemblies would have numerous applications as reactive separation media and for applications in spatially subcomparmentilizing biochemical processes. By varying the ratio of FUS LC to RGG, we observed several interesting hirearchical assemblies. We show that salt concentration affects interfacial tension and the structure of the two phases. We modulate condensate viscoelasticity by introducing a β-amyloid peptide into the RGG phase which affects miscibility of the two phases. Interestingly, we produce multiphasic condensates of FUS LC and RGG by kinetically trapping FUS LC inside RGG. In addition, to systematically coordinate the interaction of the two phases, we developed a designer surfactant protein that acts to reduce the FUS LC – RGG interfacial tension and allows complex multiphase partitioning. In sum, we demonstrate feasibility of generating novel structures by interfacing two phases that normally do not integrate. This study provides insight on the physical and chemical principles governing self-assembly of these soft peptide materials and reveals new approachs to rationally design multphase protein condensates for applications in chemistry and cellular engineering.

## Results and Discussion

### FUS LC and RGG condensates form immiscible phases modulated by salt concentration

The N-terminal QGSY-rich domain of Human RNA-binding protein FUS is characterized by low sequence complexity and mediates liquid-liquid phase separation (LLPS) of FUS (24, 26). We made condensates of the 163-a.a domain of FUS (FUS LC), by fusing two FUS LC domains together to make a FUS LC dimer, as it is known that multivalency of Intrinsically-disorderd regions (IDR) promote LLPS (23, 27). We added a protein solubility tag, maltose binding protein (MBP), followed by a HRV-3C protease cleavage site, and inserted a fluorescent protein mTagBFP2 in between 2 FUS LC domains to enable tracing of the condensates by fluorescence microscopy (Fig.S1a). Following digestion by HRV3C, phase transition occured. By preincubating MBP-FUSLC-mTagBFP2-FUSLC with HRV-3C at 42 ° before diluting into the imaging wells, the protein formed spherical droplets in vitro at micromolar level concentration (Fig.S1b). Droplets were capable of fusing while maintaining their spherical shape (Fig.S1c), indicating a liquid character. FUS LC condensates are known to behave like aging Maxwell fluids (8), and would gradually mature over time to transition into gels or fibrous aggregates (28). Fluorescence recovery after photobleaching (FRAP) experiments 2 hours after assembly of FUSLC-mTagBFP2-FUSLC droplets indicate a slow and partial recovery after photobleaching, confirming a gel-like character (Fig. S1d). Freshly prepared FUSLC-mTagBFP2-FUSLC droplets could not be dissolved by high concentration salt solution (Fig. S10), confirming its gel behavior, similar to an aged FUS condensate (29). It has been suggested that FUS LC condensates ‘salts out’ the solution (LLPS enhanced by increased NaCl concentration), indicating a significant contribution from hydrophobic interactions to LLPS (24). We did not observe the formation of aggregates or fibrils from FUSLC-mTagBFP2-FUSLC condensates in mTagBFP2 fluorescence channel over the course of 5 days (Fig. S11). In addition, excitation of mTagBFP2 by the 405 nm laser notably accelerated transition of spherical droplets into dynamically arrested gels (not shown).

FUS LC and RGG polypeptide scaffolds are immiscible and form discrete condensates when reconstituted biochemically in vitro or when expressed in yeast and mammalian cells (22). In this study, we focused on the immiscibility of the two phases, and their mesoscale structures. We first formed FUSLC-mTagBFP2-FUSLC condensates in the imaging well, then mixed them with RGG-GFP-RGG condensates. Confocal microscopy (Fig. 1a) shows that the two phases did not mix, and instead partially wetted each other, indicating similar interfacial tensions of the FUS LC and RGG phases (6). Switching the order of assembly produced similar partially wetted structures. To estimate the interfacial tensions, we measured the contact angles between the 3 phases (Fig. 1b). We obtained the values of the two angles, α = 112.0 ± 8.2 °and β = 107.5 ± 4.8°, respectively. Solving the equations *γ*_*FR*_ *+ γ*_*FW*_ cos α *+γ*_*RW*_ cos *β* = 0 and *γ*_*FW*_ sin α − γ_*RW*_ sin *β* = 0, where *γ*_*FR*_ is the interfacial tension between FUS LC and RGG phases, *γ*_*FW*_ is the interfacial tension between FUS LC phase and water, and *γ*_*RW*_ is the interfacial tension between RGG phase and water, we obtained that *γ*_*FW*_ = 1.02 *γ*_*RW*_ and *γ*_*FR*_ = 0.68 *γ*_*RW*_ . The surface tension of RGG phase (RGG-GFP-RGG) has previously been measured by micropipette aspiration at *γ*_*RW*_ = 0.159 *mN*/*m* (7). Therefore, we estimate that *γ*_*FW*_ = 0.162 *mN*/*m* and *γ*_*FR*_ = 0.108 *mN*/*m*. The estimated surface tension of FUS LC is in line with previously reported values obtained from molecular simulations (30-32). These condensates are not ideal liquids and rather show viscoelastic behavior, which is neglected in the force balances.

**Fig. 1.**
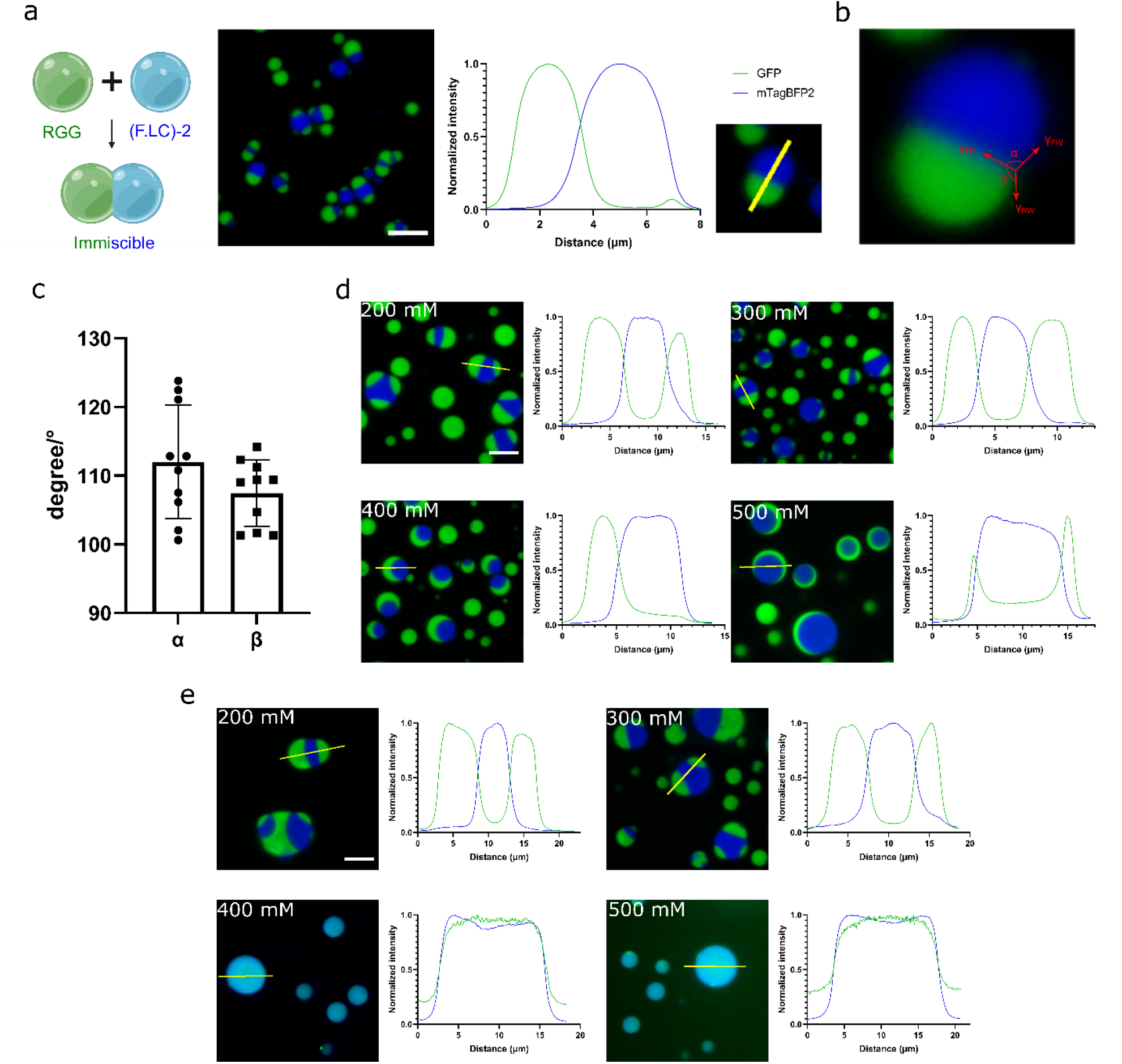
RGG and FUS LC condensates are immiscible and show salt-dependent interfacial tension. **1a**. FUSLC condensates (labeled in blue) and RGG condensates (labeled in green) are immiscible and partially-wetting. FUSLC-mTagBFP2-FUSLC droplets were pre-formed at 6 μM concentration, then 6 μM final concentration of RGG-GFP-RGG were added into the imaging well. Scale bar: 10 μm. Right, fluorescence intensity profile. **1b**. Contact angles between FUS LC phase and RGG phase. **1c**. Measured contact angles between FUS LC and RGG phase. n = 10. Error bars are standard deviations. **1d**. Interfacial tension between RGG phase and FUS LC phase is modulated by salt concentrations. MBP-FUSLC-mTagBFP2-FUSLC was incubated with HRV-3C at 42 ° before being diluted into imaging well. Then, MBP-RGG-GFP-RGG was incubated with HRV-3C at 42 ° and added into the wells containing FUS LC droplets. 5 μM final concentration of RGG-GFP-RGG and FUSLC-mTagBFP2-FUSLC final concentrations in a buffer of 20 mM HEPES and varying NaCl concentrations at pH 7.4. **1e**. 3 hr after RGG and FUS LC condensates were assembled at varying salt concentrations. At 200 mM and 300 mM NaCl concentrations, RGG and FUS LC condensates remain largely partially wetted after 3 hr. At 400 mM and 500 mM NaCl concentrations, however, RGG condensates dissolved and partitioned into the FUS LC phase.

We performed the above experiments at low salt concentrations. At higher salt concentrations (400 and 500 mM), we discovered that FUSLC-mTagBFP2-FUSLC and RGG-GFP-RGG formed a core-shell structure, with the RGG phase as the outer shell (Fig. 1d). This suggests that high salt concentration decreased the interfacial tension(s) between the RGG phase and the FUS LC phase or the outer aqueous phase by weakening the homotypic interactions between the RGG molecules. This is consistent with the salt-dependent rheological properties of the condensates (33) and complex coacervates (34). However, the RGG-GFP-RGG condensates were not stable at these salt concentrations and dissolved rather rapidly (Fig. 1e). Surprisingly, after dissolution, RGG-GFP-RGG partioned into the FUS LC condensates as well as the outer aqueous phase. We posit that weakened homotypic interactions following dissolution of the RGG-GFP-RGG condensates and the loss of the surface tension led to this co-partioning of RGG into FUS LC phase.

For complex coacervates of charged polymers, it is proposed that differences in critical salt concentrations result in immiscibility of multiphases (15). We observed such salt-dependent immiscibility change for the FUS LC – RGG multiphases (Fig. 1d, e). Although FUS LC is mostly uncharged, RGG has a realtively high charge density. The shift from immiscibility to miscibility above RGG critical salt concentrations and changes in interfacial tension with respect to salt concentration suggest the origin of FUS LC and RGG immiscibility is their charge differences. Consistent with this notion, when we used a FUS LC variant containing 32 segregated charged residues, we observed the 2 IDPs mixing (22). In addition, above the critical salt concentration, droplets dissolve and interfacial tension drops to zero (34). The loss of RGG phase along with its surface energies result in a reduced energy barrier for RGG to partition into the FUS LC phase, as predicted by theroetical studies (35, 36). Indeed, we observed that below the saturation concentration, RGG-GFP-RGG peptide from the aqueous phase partitions into the FUS LC phase (Fig. S21). Raising salt concentration favors RGG-GFP-RGG paritioning into the outer aqueous phase rather than the FUS LC phase (Fig. S21). Overall, from the drastic change to partitioning in Fig. 1d-e after dissolution of the RGG phase, we conclude that charge differences between FUS LC and RGG and their interfacial tension contribute to the immiscibility between FUS LC and RGG phases.

### Effect of condensate liquidity on miscibility

We have shown that surface properties and electrostatic interactions of the condensates play a crucial role in structuring the condensate mixture. We next ask how does condensate viscoelasticity or liquidity affect the hiearchy of assembly. To address this, we syntheized additional peptides which modulate condensate liquidity and miscibility.

To start, we wanted a pair of IDPs that form well-mixed co-condensates rather than discrete ones. For this purpose, we tested the full-length FUS protein (MBP-FUS _FL_-mCherry) with LAF-1 RGG in vitro. Unlike the mostly uncharged LC region, FUS _FL_ contains RGG domains similar to LAF-1 RGG, and co-condense with LAF-1 RGG in yeast (22). Following HRV-3C cleavage of the MBP domain, FUS _FL_ -mCherry was able to phase separate into spherical droplets indicative of their surface tension and liquid-like character. Mixing of FUS _FL_ condensates with LAF-1 RGG condensates produced well-mixed homogeneous condensates (Fig. 2a).

**Fig. 2.**
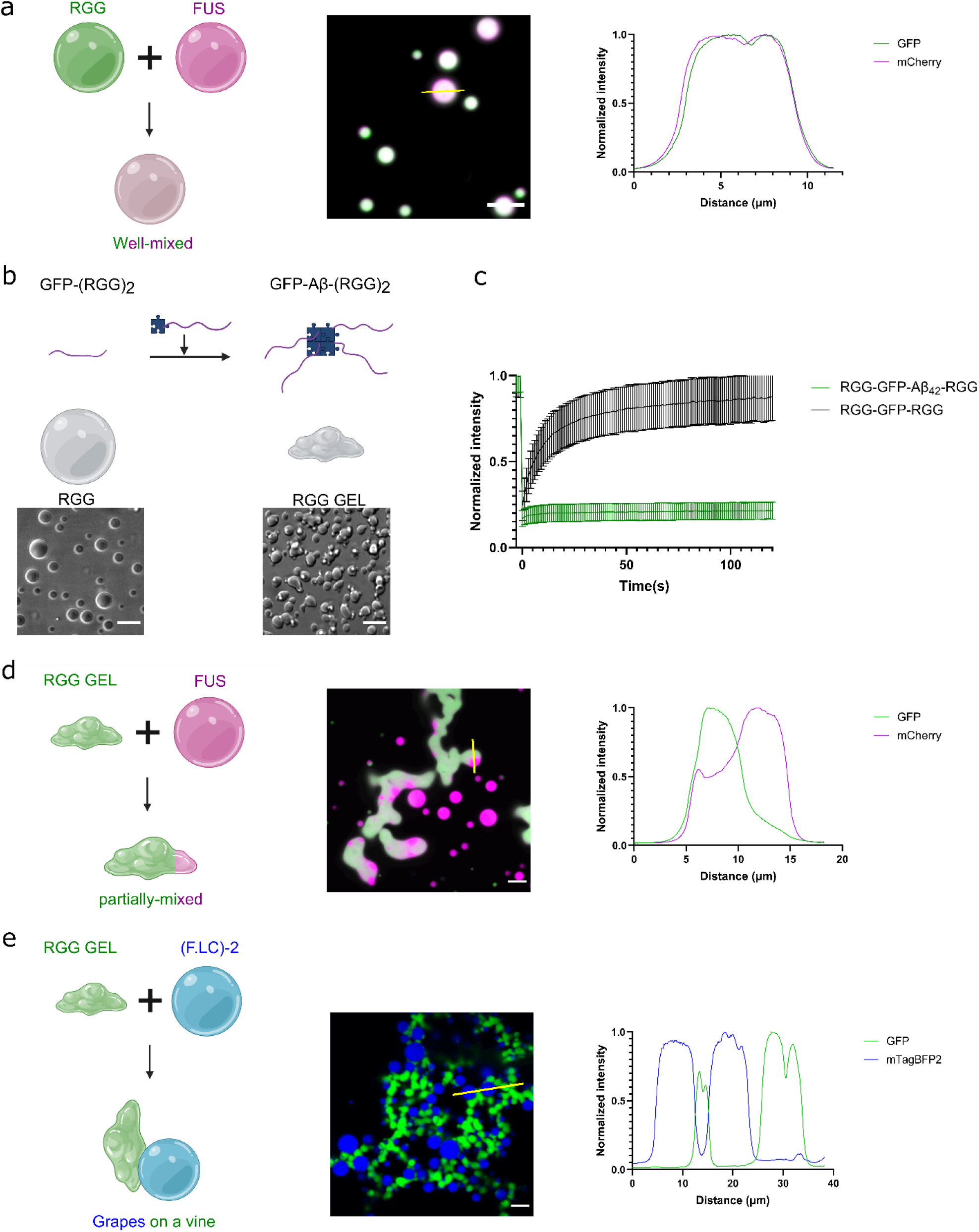
Modulating condensate miscibility by tuning viscoelasticity. **2a**. FUS _FL_ and RGG co-condense to form well-mixed condensates. 5.5 μM FUS _FL_-mCherry condensates were formed in the imaging well, then 0.9 μM RGG-GFP-RGG was added into the well. The two proteins formed co-condensates. **2b**. fusion of RGG dimer with β-amyloid (Aβ_1-42_) petpide altered condensate morphology, indicated by the increased irregularity of the Aβ containing condensates. **2c**. FRAP experiments suggest that fusion with Aβ significantly hardened the condensates, indicating a gel-like rheology. n=10. Error bars are standard deviations. **2d**. Aβ containing RGG condensates no longer well-mixed with FUS _FL_ condensates, indicating viscoelasticity affects condensates miscibility. 9 μM RGG-GFP-Aβ_42_-RGG formed condensates in the imaging well and allowed to mature into gel, then 6 μM FUS _FL_-mCherry was added. RGG-GFP-Aβ_42_-RGG and FUS _FL_-mCherry only partially mixed, with FUS _FL_-mCherry partioning in the RGG-GFP-Aβ_42_-RGG phase while RGG-GFP-Aβ_42_-RGG largely excluded in the FUS _FL_-mCherry phase. **2e**. Aβ containing RGG condensates and FUS LC condensates formed grapes on a vine structure. 9 μM RGG-GFP-Aβ_42_-RGG formed condensates in the imaging well and allowed to mature into gel, then 5 μM FUSLC-mTagBFP2-FUSLC was added. Scale bars: 10 μm.

We now had in hand a system of immiscible discrete condensates, FUSLC-mTagBFP2-FUSLC [(F.LC)_2_ for simplicity] and RGG-GFP-RGG, and a system of homogeneous co-condensates, FUS _FL_-mCherry and RGG-GFP-RGG. Next, we engineered a version of RGG condensates to reduce fluidity and increase viscoelasticity. To achieve this, we inserted an β-amyloid (Aβ_1-42_) petpide in RGG-GFP-RGG (MBP-RGG-GFP-Aβ_42_-RGG). Aβ peptide is the proteolytic product of the amyloid precursor protein and is associated with Alzheimer’s disease. The peptide is prone to the formation of amyloid fibrils, protofibrils, and oligomers (37, 38). Fusion with MBP made the recombinant protein MBP-RGG-GFP-Aβ_42_-RGG soluble, and cleavage of MBP by HRV3C released RGG-GFP-Aβ_42_-RGG and initiated formation of gel-like condensates (Fig. 2b). These Aβ_42_ containing RGG condensates are notably distinguishable from their non-Aβ containing conterparts, hallmarked by their non-spherical and irregular morphology indictive of a gel phase (Fig. 2b). FRAP experiments confirm that these condensates show gel-like behavior with no internal diffusion (Fig. 2c). In comparison, free Aβ_42_ peptide had no effect on RGG-GFP-RGG liquidity (Fig. S16). We rationalize that Aβ oligomerization may caused RGG cross-linking leading to hardened gel-like condensates. For simplicity, we named this condensate model RGG _gel_ .

To probe how does viscoelasticity affect condensate miscibility, we tested mixing RGG _gel_ condensates with FUS based condensates. Unlike RGG and FUS _FL_ which formed co-condensates, when we allowed RGG _gel_ to mature into gels then added FUS _FL_, RGG _gel_ condensates and FUS _FL_ condensates were partially immiscible (Fig. 2d). This large change in partitioning indicates that condensate liquidity controls miscibility of the disordered scaffold proteins. This is potentially caused by the increased elasticity and reduced internal diffusion of RGG _gel_. To overcome this miscibility barrier caused by increased viscoelasticity, we manually increased entropy of the system by premixing the two components together before forming the condensates (Fig. S17). Since mixing happened before RGG _gel_ could mature into gels, the condensates were well-mixed. Similar to (F.LC)_2_ and RGG condensates which formed immiscible phases, (F.LC)_2_ and RGG _gel_ also showed immiscibility, and formed ‘grapes on a vine’ structure (Fig. 2e). Based on the transition in miscibility from Fig. 2a to 2d, we conclude that IDP mutations that promote liquid-gel transitions can impact IDP miscibility, a result that has important implications for the formation of multiphase condensates

### Kinetically driven, multiphase structuring of condensates

Dynamical out-of-equilibrium transitions have been shown to generate multiphase double-emulsions. For example, reentrant phase transition induced by RNA led to vacuole substructure formation in synthetic peptide and IDP-based condensates (17). Similar structures were found in RNA-PEG system (18). These structures form due to a rapid composition change leading to the system deviating from thermodynamic equilibrium (18). Moreover, hollow vacuole substructures found in living cells indicate the potential involvement of such kinetic processes in cellular organization (39, 40). Besides the implication in life science, emusions broaden the range of assemblies that can be built from immiscible phases and find applications in various manufacturing processes (41).

We observed kinetically driven double-emulsion condensates that form in the absence of RNA, requiring only protein components (Fig. 3a). We wanted to further explore use of this process to generate multiphase droplets. We fused RGG-GFP-RGG with an N-terminal MBP, followed by an HRV-3C protease cut site. The resulting protein, MBP-RGG-GFP-RGG, was co-incubated with MBP-FUSLC-mTagBFP2-FUSLC and HRV-3C at 42°, before being diluted into the imaging well for confocal imaging. When co-incubated at an excess of MBP-RGG-GFP-RGG compared to MBP-FUSLC-mTagBFP2-FUSLC, kinetically trapped FUSLC-mTagBFP2-FUSLC droplets formed inside the RGG-GFP-RGG droplets (Fig. 3b), forming kinetically trapped multiphase structures. However, when co-incubated at an excess of MBP-FUSLC-mTagBFP2-FUSLC relative to MBP-RGG-GFP-RGG, no kinetically trapped droplets were formed, and the two phases form distinct structures dependent on the molar ratio of the two proteins (Fig. 3c,d).

**Fig. 3.**
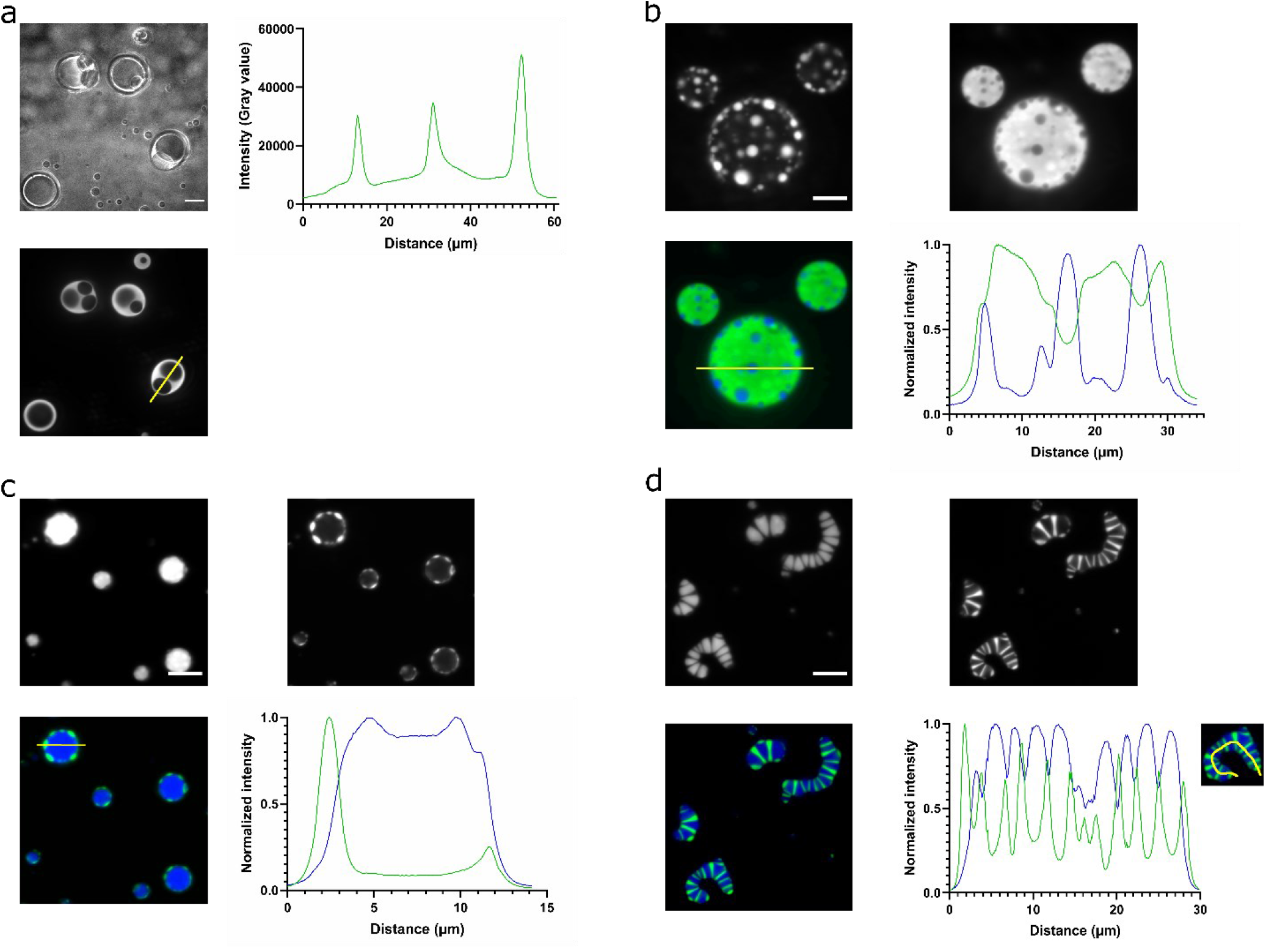
Producing multiphase condensates through kinetic trapping. **3a**. Formation of RGG double emulsion droplets. Upper left, DIC image; lower left, GFP fluorescence image; upper right, GFP fluorescence intensity profile. 10 μl 60 μM MBP-RGG-GFP-RGG and 0.5 μl 1 mg/ml HRV-3C was co-incubated at 42 ° for 30 s then 5 μl of the mixture was diluted 6-fold in 20 mM HEPES buffer, PH 7.4 for imaging. Scale bar: 20 μm. **3b**. Kinetically trapping FUS LC phase inside RGG phase. Co-incubation of MBP-FUSLC-mTagBFP2-FUSLC and MBP-RGG-GFP-RGG with HRV-3C at 42 ° at a ratio of 1:2.5 before being diluted into imaging well. Upper left, fluorescence channel for mTagBFP2; upper right, fluorescence channel for GFP; lower left, overlay; lower right, fluorescence intensity profile. 6 μM FUSLC-mTagBFP2-FUSLC and 15 μM RGG-GFP-RGG final concentration. Scale bar: 10 μm. **3c**. Formation of ‘golf-ball’ structures. Co-incubation of MBP-FUSLC-mTagBFP2-FUSLC and MBP-RGG-GFP-RGG with HRV-3C at 42 ° at a ratio of 2:1 before being diluted into imaging well. Upper left, fluorescence channel for mTagBFP2; upper right, fluorescence channel for GFP; lower left, overlay; lower right, fluorescence intensity profile. 8 μM FUSLC-mTagBFP2-FUSLC and 4 μM RGG-GFP-RGG final concentration. Scale bar: 10 μm. **3d**. Formation of ‘centipede’ structures. Co-incubation of MBP-FUSLC-mTagBFP2-FUSLC and MBP-RGG-GFP-RGG with HRV-3C at 42 ° at a ratio of 3:1 before being diluted into imaging well. The two phases formed a ‘centipede’ like structure. Upper left, fluorescence channel for mTagBFP2; upper right, fluorescence channel for GFP; lower left, overlay; lower right, fluorescence intensity profile. 6 μM FUSLC-mTagBFP2-FUSLC and 2 μM RGG-GFP-RGG final concentration. Scale bar: 10 μm.

It is theorized that kinetically trapped hollow droplets form as a result of rapid composition change leading to an out-of-equilibrium state deviating from the binodal (18). This was corroborated from our observation on the FUS _FL_-mCherry and RGG-GFP-RGG mixtures (Fig.S13-S15). When we reduced the composition change rate, formation of the kinetically trapped droplets was prevented (Fig. 2a, right). Similarly, controlling MBP-RGG-GFP-RGG cleavage rate and molar ratio of proteins yielded different structures. These results suggest a kinetic aspect in the process of condensate assembly, namely, composition change per time affects the final sturcture. These different structures under different formation rate and stoichiometry suggest a kinetically driven out-of-equillibrium formation of the condensate mixture that is history-dependent.

### Multiphase assemblies of FUS LC and RGG dictated selectively by surfactant type

We have shown that FUS LC and RGG droplets form immiscible phases that partially contact each other. This is due to the similar interfacial tensions between the RGG, FUS LC, and aqueous phases. We next sought to reduce the interfacial tension between FUS LC and RGG phases by using a designer surfactant protein and build a multiphase condensate (Fig. 4a). Such a strategy would be needed for example inside living cells where variation of salt concentration or kinetic trapping would be much less feasible. We first tested some commercial amphiphilic surfactant molecules, which were wholly ineffective and did not alter the partially wetted configuration of the immiscible phases (Fig. S19). We also tested the effect of RNA on the structures of the condensates, as previous report shows formation of vacuoles as a result of RNA induced reentrant phase transition (17). We observed reentrant transition after polyU RNA initially dissoloving FUS LC and RGG condensates (Fig. S20). Nevertheless, FUS LC and RGG condensates remained demixed, and no core-shell sturctures were formed (Fig. S20).

**Fig. 4.**
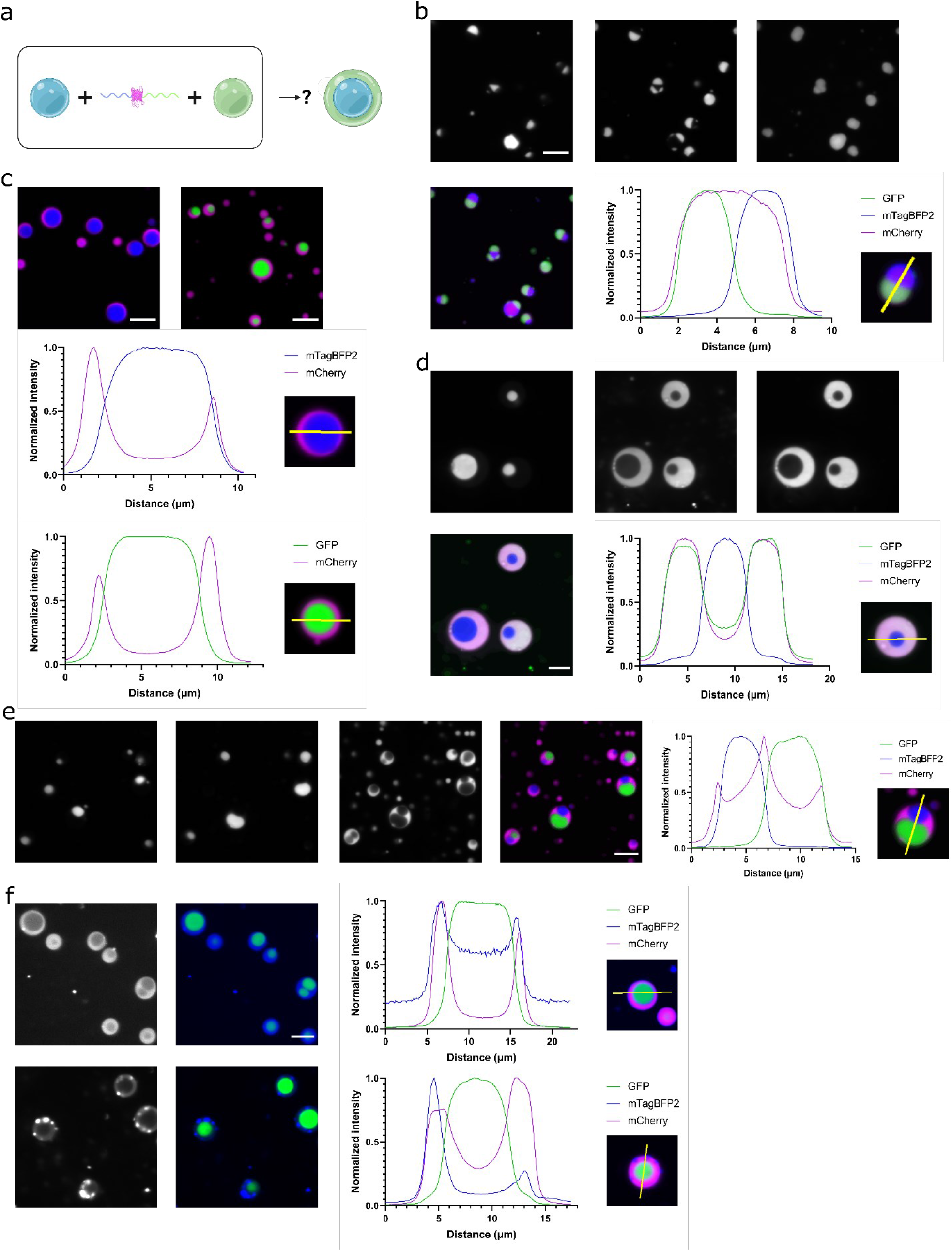
Building multiphasic condensates using designer surfactant proteins. **4a**. Design of recombinant surfactant protein branching FUS LC phase and RGG phases. **4b**. FUSLC-mCherry-RGG partions into both FUS LC and RGG phases. Top panels from left to right: FUSLC-mTagBFP-FUSLC phase, RGG-GFP-RGG phase, FUSLC-mCherry-RGG fluorescence; bottom panels from left to right: overlay of fluorescence, fluorescence intensity profile. 1 μM final concentration of FUSLC-mCherry-RGG was added into well containing 2 μM FUSLC-mTagBFP2-FUSLC and 7 μM RGG-GFP-RGG. Scale bar: 10 μm. **4c**. FUSLC-mCherry-RGG partitions to the surfaces of FUSLC-mTagBFP2-FUSLC condensates (left) and RGG-GFP-RGG condensates (right). Blue, FUSLC-mTagBFP2-FUSLC; green, RGG-GFP-RGG; magenta, FUSLC-mCherry-RGG. Scale bar: 10 μm. **4d**. Core-shelled multiphase condensate. Top panels from to right: FUSLC-mTagBFP2-FUSLC condensates as inner core, RGG-GFP-RGG as outer shell, FUSLC-mCherry-RGG as surfactant; bottom panels from left to right: overlay of fluorescence, fluorescence intensity profile. Condensates of 1 μM FUSLC-mTagBFP2-FUSLC was preformed, 1 μM FUSLC-mCherry-RGG was then added and 5.5 μM RGG-GFP-RGG was last added. Scale bar: 10 μm. **4e**. Dual-cored multiphase condensates. Left to right: FUSLC-mTagBFP2-FUSLC, RGG-GFP-RGG, FUSLC-mCherry-RGG as outer shell, overlay of fluorescence, intensity profile. 3 μM FUSLC-mTagBFP2-FUSLC and 3 μM RGG-GFP-RGG were pre-assembled, then 1 μM FUSLC-mCherry-RGG was added. Scale bar: 10 μm. **4f**. FUSLC-mTagBFP2-FUSLC as the outer shell in a multiphase structure. Top panel: 3 μM RGG-GFP-RGG, 2 μM FUSLC-GST-mCherry-RGG, and 3 μM FUSLC-mTagBFP2-FUSLC. Lower panel, more FUSLC-mTagBFP2-FUSLC added into well, final concentration 6 μM. Left to right: FUSLC-mTagBFP2-FUSLC fluorescence, overlay of RGG-GFP-RGG fluorescence with FUSLC-mTagBFP2-FUSLC fluorescence (mCherry fluorescence not shown for ease of observation, fluorescence intensity profile. Scale bar: 10 μm.

Next, we set out to rationally build synthetic biopolymer condensates using protein blocks. Previous efforts were made to build a surfactant protein that localizes at and recruits cargo to the RGG condensate-water interface (19, 42). This was achieved by fusion of a hydrophilic protein domain Glutathione S-transferase (GST) with a condensate domain RGG, which results in an amphiphilic protein that stabilizes the RGG condensate-water interface. In our initial attempt to bridge FUS LC and RGG, we designed an amphiiphile-like protein that contains a FUS LC domain and an RGG domain, with a fluorescent reporter mCherry in between. We reasoned that since FUS LC and RGG generally do not interact, the resulting protein, FUSLC-mCherry-RGG, would preferentially stay at the interface of FUS LC – RGG condensates. However, confocal imaging shows that the protein partitioned homogeneously into both RGG and FUS LC phases, and hence was not surface-active (Fig. 4b). Although FUSLC-mCherry-RGG could partition into both phases, FUS LC and RGG phases remained demixed. To make a surface-active protein that could act as a surfactant between FUS LC and RGG phases, we added a hydrophilic GST domain between FUSLC and RGG, to make FUSLC-GST-mCherry-RGG; the increased hydrophilicity excludes the surfactant from the condensed phases. We found that adding FUSLC-GST-mCherry-RGG into wells containing FUSLC-mTagBFP2-FUSLC droplets or RGG-GFP-RGG droplets resulted in surface localization of the surfactant at both condensates (Fig. 4c). When we first formed FUSLC-mTagBFP2-FUSLC condensates, then added the surfactant FUSLC-GST-mCherry-RGG, and lastly added RGG-GFP-RGG, we formed core-shelled structured, multiphasic condensates with FUSLC as the inner core and RGG as the outer shell (Fig. 4d). The presence of surfactant protein reduced interfacial tension between FUS LC and RGG and induced RGG-GFP-RGG to wetting the interface of FUS LC condensates (Fig. 4d).

As kinetics affect structuring of condensates, we wondered whether under specific conditions we could form triphase condensates. At sufficiently high concentrations, the designer surfactant protein FUSLC-GST-mCherry-RGG could phase separate to form a distinct phase on its own. Thus, we decided to vary stochiometry and the order of component addition. We first formed immiscible FUSLC-mTagBFP2-FUSLC and RGG-GFP-RGG condensates as in Fig. 1a, then added FUSLC-GST-mCherry-RGG. This change of order of addition resulted in a condensate structure with three distinct phases (Fig. 4e). The surfactant acted as an outer shell enveloping FUSLC and RGG phases, and led to a hetero-core-shell structure with both the FUS LC and RGG phases as the inner core and the surfactant as the outer shell. Dual core condensates with FUS LC and RGG as inner core can be observed through confocal imaging. This is likely the result of the hydrophilic contribution from GST to the surfactant that made it a bi-functional surfactant capable of staying at both FUS LC and RGG surfaces (Fig. 4c). The difference in multiphasic condensate structure caused by different order of component addition highlight that the formation of multiphasic structure is history-dependent and the system likely assumed a local free energy minimum. It is known that emulsion systems do not always assume the lowest energy state predicted by thermodynamic analysis, rather kinetic factors affect their structure (43). This agrees with the kinetically stable but thermodynamically unstable nature of emulsion systems (41).

Lastly, we asked whether we could reverse the order and make RGG-GFP-RGG the inner core, and FUSLC-mTagBFP2-FUSLC the outer shell in a multiphasic condensate. RGG-GFP-RGG condensates were first formed, then the surfactant was added, and last FUSLC-mTagBFP2-FUSLC was added (Fig. 4f). FUSLC-mTagBFP2-FUSLC weakly wetted the RGG-GFP-RGG interface. However, there is an elevated concentration of FUSLC-mTagBFP2-FUSLC in the RGG-GFP-RGG inner core, likely due to the higher interfacial tension of FUS LC than RGG with water leading to instability of the system. Adding more FUSLC-mTagBFP-FUSLC into the well led to its self-coacervation into smaller FUSLC droplets near the interface. This indicates that it is energetically costly for FUS LC to wet at the interface which is larger in surface area because of its higher surface tension.

Overall, through the use of rationally designed surfactant proteins containing IDRs from two partially wetting condensed phases, we effectively reduced the interfacial tension between the two phases and assembled multiphase structures from minimalistic condensate models. This result highlights that engineering condensate multiphasic structure is feasible via adjusting interfacial tension, and illustrates physical principles guiding their formation and engineering.

## Conclusions

In this study, we built a model FUS LC condensate system whose formation can be controlled by adding a protease HRV-3C. FUS LC condensates are spherical and able to fuse, indicating they are partially liquid. FRAP on FUS LC condensates reveal that they have slow internal diffusivity and are gel-like. FUS LC condensates and RGG condensates are immiscible in vitro, resulting in partial wetting with each other. Various structures can be formed out of the two phases under different conditions. We modulated condensate viscoelasticity which affected condensate miscibility. We show that we could make FUS LC condensates kinetically trapped inside the RGG condensates, resulting in multiphase structures. In addition, we could also build multiphasic FUS LC / RGG condensates by using a designer surfactant protein that branches the two proteinaceous phases and thus reducing the interfacial tension. The structure of the multiphasic condensates is history-dependent thereby showing a kinetic aspect of their assembly. Through this study, we hope to provide insight into principles guiding the self-assembly of complex coacervate soft matter and a strategy for rationally designing multiphasic condensates from immiscible phases using the generic surfactant approach.

## Acknowledgements

This work was funded by the NSF MRSEC Grant (DMR-2309034). The authors would like to thank Cell & Developmental Biology (CDB) Microscopy Core, Jasmine Zhao and Andrea Stout at the University of Pennsylvania for providing the facility for FRAP and for technical support.

## Author contributions

M.G, D.A.H. and M.C.G. designed the experiments. M.G. performed the experiments. M.G, D.A.H. and

M.C.G. wrote the manuscript.

## Data availability

Data available on request from the authors.

### Methods

### Cloning

The 163 amino acid long QGSY-rich domain of FUS (referred to FUS LC) was amplified by PCR and inserted into an empty pET vector by In-Fusion cloning (Takara Bio, Kusatsu, Shiga, Japan). A second FUS LC domain was inserted by restriction cloning to create FUSLC-FUSLC. A plasmid encoding mTagBFP2 (44) was a gift from Michael Davidson (Addgene plasmid # 54572 ; http://n2t.net/addgene:54572 ; UnderRRID:Addgene_54572). MBP with a HRV-3C protease cleavage site and linker and mTagBFP2 were inserted by In-Fusion cloning to create MBP-FUSLC-mTagBFP2-FUSLC. Other proteins were cloned in a similar manner. The plasmids encode aminoglycoside phosophotransferase that confers resistance to kanamycin. All primers were ordered from Integrated DNA Technologies (Coralville, IA). Plasmids were sequence confirmed by Next-Gen Sequencing (GENEWIZ, South Plainfield, NJ).

### Protein purification

All MBP tagged proteins were purified by Amylose resin (New England Biolabs, Ipswich, MA). Proteins without MBP tags were tagged with 6-His and purified by HisPur Ni-NTA resin (Thermo Scientific, Waltham, MA). Plasmids containing the encoded proteins were transformed into BL 21 (DE3) competent cells. Starter cultures were grown in 5 mL Luria Broth supplemented with 50 μg/mL kanamycin overnight at 37 °C. The cultures were then innoculated into 500 mL Terrific Broth – Autoinduction media with 50 μg/mL kanamycin at 37 °C to an optical density of 0.5-1.0 then grown at 18 °C overnight. Cells were pelleted by centrifugation at 5000 g for 10 min and stored at -20 °C. Pellets were resuspended in buffer A (20 mM HEPES, 500 mM NaCl, PH 7.4) supplemented with 20 mM imidazole (for proteins purified by Ni-NTA resins), protease inhibitor (EDTA-free, Roche, Basel, Switzerland), 0.1% CHAPS, 1 mg/mL lysozyme and lysed by sonification. Lysates were centrifuged at top speed (15,000 g) for 20 min at room temperature for MBP tagged proteins and at 42 °C for non-MBP tagged phase separating protein (RGG-GFP-RGG). Supernatant was incubated with the corresponding resins, washed by buffer A (supplemented with 20 mM imidazole for proteins purified by Ni-NTA resins), and eluted on a disposable 5 mL polypropylene column (Thermo Scientific). Elution buffer for amylose resins is buffer A with 50 mM maltose, and for Ni-NTA resins buffer A with 500 mM imidazole. Protein eluted in high imidazole buffer (RGG-GFP-RGG) was dialyzed into imidazole-free buffer (buffer A) at 42 °C. Protein concentrations were determined by NanoDrop ND-1000 spectrophotometer (Thermo Fisher Scientific). Protein solutions were flash frozen in liquid nitrogen and stored at -80 °C.

### Protein gel electrophoresis

Protein samples were denatured by NuPAGE (Invitrogen, Carlsbad, CA) LDS Sample Buffer and reduced by NuPAGE Sample Reducing Agent, and incubated at 98 °C for 3 min. Samples and Novex Sharp Pre-stained Protein Standard (Invitrogen, Carlsbad, CA) were loaded on NuPAGE 4-12% Bis-Tris gels in a XCell SureLock Electrophoresis System (Invitrogen) at 200 V for 35 min. Gels were stained by SimplyBlue and imaged by a LI-COR (Lincoln, NE) Odyssey M imaging system.

### Confocal microscopy

Proteins were thawed at 42 °C for RGG-GFP-RGG and at room temperature for MBP tagged proteins, and centrifuged at 14,000 g for 3 min to remove aggregates. MBP tagged proteins were incubated with HRV-3C protease (Sigma-Aldrich, St. Louis, MO) at 42 °C to remove the MBP tag while preventing phase separation. Phase separation was induced by diluting down the salt concentration of the protein solution using salt-free buffer (20 mM HEPES, PH 7.4). Protein samples were placed in CultureWell chambered coverglass (Invitrogen, Carlsbad, CA) and imaged on a Leica (Wetzlar, Germany) DMi8 inverted microscope equipped with a confoal spinning disk unit (Spectral Applied Research, Richmond Hill, Canada), with a 63x 1.4 NA plan-apochromatic oil-immersion objective and a Hamamatsu ORCA-Flash 4.0 V2 digital sCMOS camera (Shizuoka, Japan). To prevent wetting of condensates at coverglass surface, imaging wells were pre-coated with 5% Pluronic F-127 (Sigma-Aldrich) solution for at least 1 hr and rinsed before use.

### Fluorescence recovery after photoblaching

FRAP experiments on FUSLC-mTagBFP2-FUSLC were performed on a Leica DMi8 inverted microscope (Leica, Wetzlar, Germany) equipped with a Leica Stellaris confocal system, and extended tunable WLL excitation from 485-790 nm plus fixed 405 nm laser. mTagBFP2 fluorescence was bleached by the 405 nm laser at 100% intensity and imaged at 1% intensity using a 63x 1.4 NA oil-immersion objective. Circular regions of 1 μm diameter were bleached in the center of protein droplets and recovery was monitored for 180 s post-bleach in the LAS X software (Leica, Wetzlar, Germany). FRAP on GFP-containing proteins were performed on the equipment previously described (42). Circular regions of ∼1 μm diameter were bleached in the center of protein droplets and recovery was monitored for around 120 s in MetaMorph software (Molecular Devices, Downingtown, PA).

