## Supplementary Materials PDF for "Assembly of hierarchical multiphase condensates using designer surfactant proteins"

**Table S1. Amino acid sequences of proteins used in this study.**

| Name | Sequence |
|---|---|
| MBP-FUSLC-mTagBFP2-FUSLC | MKIEEGKLVIWINGDKGYNGLAEVGKKFEKDTGIKVTVEHPDKLEEKFPQVA<br>ATGDGPDIIFWAHDRFGGYAQSGLLAEITPDKAFQDKLYPFTWDAVRYNGK<br>LIAYPIAVEALSLIYNKDLLPNPPKTWEEIPALDKELKAKGKSALMFNLQEPYF<br>TWPLIAADGGYAFKYENGKYDIKDVGVDNAGAKAGLTFLVDLIKNKHMNA<br>DTDYSIAEAAFNKGETAMTINGPWAWSNIDTSKVNYGVTVLPTFKGQPSKP<br>FVGVLSAGINAASPNKELAKEFLENYLLTDEGLEAVNKDKPLGAVALKSYEEE<br>LVKDPRIAATMENAQKGEIMPNIPQMSAFWYAVRTAVINAASGRQTVDEA<br>LKDAQTGGGGSGGGGSLEVLFQGPGHMASNDYTQQATQSYGAYPTQPG<br>QGYSQQSSQPYGQQSYSGYSQSTDTSGYGQSSYSSYGQSQNTGYGTQSTP<br>QGYGSTGGYGSSQSSQSSYGQQSSYPGYGQQPAPSSTSGSYGSSSQSSSYG<br>QPQSGSYSQQPSYGGQQQSYGQQQSYNPPQGYGQQNQYNSGSGGMVS<br>KGEELIKENMHMKLYMEGTVDNHHFKCTSEGEGKPYEGTQTMRIKVVEGG<br>PLPFAFDILATSFLYGSKTFINHTQGIPDFFKQSFPEGFTWERVTTYEDGGVLT<br>ATQDTSLQDGCLIYNVKIRGVNFTSNGPVMQKKTLGWEAFTETLYPADGGL<br>EGRNDMALKLVGGSHLIANAKTTYRSKKPAKNLKMPGVYYVDYRLERIKEA<br>NNETYVEQHEVAVARYCDLPSKLGHKLNGGSGSMASNDYTQQATQSYGAY<br>PTQPGQGYSQQSSQPYGQQSYSGYSQSTDTSGYGQSSYSSYGQSQNTGYG<br>TQSTPQGYGSTGGYGSSQSSQSSYGQQSSYPGYGQQPAPSSTSGSYGSSSQ<br>SSSYGQPQSGSYSQQPSYGGQQQSYGQQQSYNPPQGYGQQNQYNSAAA<br>LEHHHHHH |
| MBP-FUS $_{FL}$-mCherry (carries a G156E mutation) | MKIEEGKLVIWINGDKGYNGLAEVGKKFEKDTGIKVTVEHPDKLEEKFPQVA<br>ATGDGPDIIFWAHDRFGGYAQSGLLAEITPDKAFQDKLYPFTWDAVRYNGK<br>LIAYPIAVEALSLIYNKDLLPNPPKTWEEIPALDKELKAKGKSALMFNLQEPYF<br>TWPLIAADGGYAFKYENGKYDIKDVGVDNAGAKAGLTFLVDLIKNKHMNA<br>DTDYSIAEAAFNKGETAMTINGPWAWSNIDTSKVNYGVTVLPTFKGQPSKP<br>FVGVLSAGINAASPNKELAKEFLENYLLTDEGLEAVNKDKPLGAVALKSYEEE<br>LVKDPRIAATMENAQKGEIMPNIPQMSAFWYAVRTAVINAASGRQTVDEA<br>LKDAQTGGGGSGGGGSLEVLFQGPGMASNDYTQQATQSYGAYPTQPGQ<br>GYSQQSSQPYGQQSYSGYSQSTDTSGYGQSSYSSYGQSQNTGYGTQSTPQ<br>GYGSTGGYGSSQSSQSSYGQQSSYPGYGQQPAPSSTSGSYGSSSQSSSYGQ<br>PQSGSYSQQPSYGGQQQSYGQQQSYNPPQGYEQQNQYNSSSGGGGGG<br>GGGGNYGQDQSSMSSGGGSGGGYGNQDQSGGGGSGGYGQQDRGGRG<br>RGGSGGGGGGGGGGYNRSSGGYEPRGRGGGRGGRGGMGGSDRGGFN<br>KFGGPRDQGSRHDSEQDNSDNNTIFVQGLGENVTIESVADYFKQIGIIKTNK<br>KTGQPMINLYTDRETGKLKGEATVSFDDPPSAKAAIDWFDGKEFSGNPIKVS<br>FATRRADFNRGGGNGRGGRGRGGPMGRGGYGGGGSGGGGRGGFPSGG<br>GGGGGQQRAGDWKCPNPTCENMNFSWRNECNQCKAPKPDGPGGGPG<br>GSHMGGNYGDDRRGGRGGYDRGGYRGRGGDRGGFRGGRGGGDRGGF<br>GPGKMDSRGEHRQDRRERPYGGSGMVSKGEEDNMAIIKEFMRFKVHME<br>GSVNGHEFEIEGEGEGRPYEGTQTAKLKVTKGGPLPFAWDILSPQFMYGSK<br>AYVKHPADIPDYLKLSFPEGFKWERVMNFEDGGVVTVTQDSSLQDGEFIYK<br>VKLRGTNFPSDGPVMQKKTMGWEASSERMYPEDGALKGEIKQRLKLKDG<br>GHYDAEVKTTYKAKKPVQLPGAYNVNIKLDITSHNEDYTIVEQYERAEGRHS<br>TGGMDELYKSSHHHHHH |

| | |
|---|---|
| RGG-GFP-RGG | MESNQSNNGGSGNAALNRGGRYVPPHLRGGDGGAAAAASAGGDDRRG<br>GAGGGGYRRGGGNSGGGGGGGYDRGYNDNRDDRDNRGGSGGYGRDR<br>NYEDRGYNGGGGGGGNRGYNNNRGGGGGGYNRQDRGDGGSSNFSRG<br>GYNNRDEGSDNRGSGRSYNNDRRDNGGDGEFLVPRGSMVSKGEELFTGV<br>VPILVELDGDVNGHKFSVSGEGEGDATYGKLTLKFICTTGKLPVPWPTLVTTL<br>TYGVQCFSRYPDHMKQHDFFKSAMPEGYVQERTIFFKDDGNYKTRAEVKF<br>EGDTLVNRIELKGIDFKEDGNILGHKLEYNYNSHNVYIMADKQKNGIKVNFK<br>IRHNIEDGSVQLADHYQQNTPIGDGPVLLPDNHYLSTQSALSKDPNEKRDH<br>MVLLEFVTAAGITLGMDELYKEFLVPRGSMESNQSNNGGSGNAALNRGGR<br>YVPPHLRGGDGGAAAAASAGGDDRRGGAGGGGYRRGGGNSGGGGGGG<br>YDRGYNDNRDDRDNRGGSGGYGRDRNYEDRGYNGGGGGGGNRGYNN<br>NRGGGGGGYNRQDRGDGGSSNFSRGGYNNRDEGSDNRGSGRSYNNDR<br>RDNGGDGLEHHHHHH |
| MBP-RGG-GFP-A$\beta_{42}$-RGG | MHHHHHHGGGSMKIEEGKLVIWINGDKGYNGLAEVGKKFEKDTGIKVTVE<br>HPDKLEEKFPQVAATGDGPDIIFWAHDRFGGYAQSGLLAEITPDKAFQDKLY<br>PFTWDAVRYNGKLIAYPIAVEALSLIYNKDLLPNPPKTWEEIPALDKELKAKG<br>KSALMFNLQEPYFTWPLIAADGGYAFKYENGKYDIKDVGVDNAGAKAGLTF<br>LVDLIKNKHMNADTDYSIAEAAFNKGETAMTINGPWAWSNIDTSKVNYGV<br>TVLPTFKGQPSKPFVGVLSAGINAASPNKELAKEFLENYLLTDEGLEAVNKDK<br>PLGAVALKSYEEELVKDPRIAATMENAQKGEIMPNIPQMSAFWYAVRTAVI<br>NAASGRQTVDEALKDAQTGGGGSGGGGSLEVLFQGPGMESNQSNNGGS<br>GNAALNRGGRYVPPHLRGGDGGAAAAASAGGDDRRGGAGGGGYRRGG<br>GNSGGGGGGGGYDRGYNDNRDDRDNRGGSGGYGRDRNYEDRGYNGGG<br>GGGGNRGYNNNRGGGGGGYNRQDRGDGGSSNFSRGGYNNRDEGSDNR<br>GSGRSYNNDRRDNGGDGEFVSKGEELFTGVVPILVELDGDVNGHKFSVSGE<br>GEGDATYGKLTLKFICTTGKLPVPWPTLVTTLTYGVQCFSRYPDHMKQHDFF<br>KSAMPEGYVQERTIFFKDDGNYKTRAEVKFEGDTLVNRIELKGIDFKEDGNIL<br>GHKLEYNYNSHNVYIMADKQKNGIKVNFKIRHNIEDGSVQLADHYQQNTPI<br>GDGPVLLPDNHYLSTQSKLSKDPNEKRDHMVLLEFVTAAGITLGMDELYKG<br>GGSDAEFRHDSGYEVHHQKLVFFAEDVGSNKGAIIGLMVGGVVIAEFLVPR<br>GSMESNQSNNGGSGNAALNRGGRYVPPHLRGGDGGAAAAASAGGDDR<br>RGGAGGGGYRRGGGNSGGGGGGGYDRGYNDNRDDRDNRGGSGGYGR<br>DRNYEDRGYNGGGGGGGNRGYNNNRGGGGGGYNRQDRGDGGSSNFSR<br>GGYNNRDEGSDNRGSGRSYNNDRRDNGGDGLEHHHHHH |
| MBP-RGG-GFP-RGG (HRV-3C cut site) | MHHHHHHGGGSMKIEEGKLVIWINGDKGYNGLAEVGKKFEKDTGIKVTVE<br>HPDKLEEKFPQVAATGDGPDIIFWAHDRFGGYAQSGLLAEITPDKAFQDKLY<br>PFTWDAVRYNGKLIAYPIAVEALSLIYNKDLLPNPPKTWEEIPALDKELKAKG<br>KSALMFNLQEPYFTWPLIAADGGYAFKYENGKYDIKDVGVDNAGAKAGLTF<br>LVDLIKNKHMNADTDYSIAEAAFNKGETAMTINGPWAWSNIDTSKVNYGV<br>TVLPTFKGQPSKPFVGVLSAGINAASPNKELAKEFLENYLLTDEGLEAVNKDK<br>PLGAVALKSYEEELVKDPRIAATMENAQKGEIMPNIPQMSAFWYAVRTAVI<br>NAASGRQTVDEALKDAQTGGGGSGGGGSLEVLFQGPGMESNQSNNGGS<br>GNAALNRGGRYVPPHLRGGDGGAAAAASAGGDDRRGGAGGGGYRRGG<br>GNSGGGGGGGGYDRGYNDNRDDRDNRGGSGGYGRDRNYEDRGYNGGG<br>GGGGNRGYNNNRGGGGGGYNRQDRGDGGSSNFSRGGYNNRDEGSDNR<br>GSGRSYNNDRRDNGGDGEFVSKGEELFTGVVPILVELDGDVNGHKFSVSGE<br>GEGDATYGKLTLKFICTTGKLPVPWPTLVTTLTYGVQCFSRYPDHMKQHDFF<br>KSAMPEGYVQERTIFFKDDGNYKTRAEVKFEGDTLVNRIELKGIDFKEDGNIL |

| | |
|---|---|
| | GHKLEYNYNSHNVYIMADKQKNGIKVNFKIRHNIEDGSVQLADHYQQNTPI<br>GDGPVLLPDNHYLSTQSKLSKDPNEKRDHMVLLEFVTAAGITLGMDELYKEF<br>LVPRGSMESNQSNNGGSGNAALNRGGRYVPPHLRGGDGGAAAAASAGG<br>DDRRGGAGGGGYRRGGGNSGGGGGGGYDRGYNDNRDDRDNRGGSGG<br>YGRDRNYEDRGYNGGGGGGGNRGYNNNRGGGGGGYNRQDRGDGGSS<br>NFSRGGYNNRDEGSDNRGSGRSYNNDRRDNGGDGLEHHHHHH |
| MBP-RGG-GFP-RGG (TEV cut site) | MKIEEGKLVIWINGDKGYNGLAEVGKKFEKDTGIKVTVEHPDKLEEKFPQVA<br>ATGDGPDIIFWAHDRFGGYAQSGLLAEITPDKAFQDKLYPFTWDAVRYNGK<br>LIAYPIAVEALSLIYNKDLLPNPPKTWEEIPALDKELKAKGKSALMFNLQEPYF<br>TWPLIAADGGYAFKYENGKYDIKDVGVDNAGAKAGLTFLVDLIKNKHMNA<br>DTDYSIAEAAFNKGETAMTINGPWAWSNIDTSKVNYGVTVLPTFKGQPSKP<br>FVGVLSAGINAASPNKELAKEFLENYLLTDEGLEAVNKDKPLGAVALKSYEEE<br>LVKDPRIAATMENAQKGEIMPNIPQMSAFWYAVRTAVINAASGRQTVDEA<br>LKDAQTNSSSNNNNNNNNNNLGENLYFQGGMESNQSNNGGSGNAALN<br>RGGRYVPPHLRGGDGGAAAAASAGGDDRRGGAGGGGYRRGGGNSGGG<br>GGGGYDRGYNDNRDDRDNRGGSGGYGRDRNYEDRGYNGGGGGGGNR<br>GYNNNRGGGGGGYNRQDRGDGGSSNFSRGGYNNRDEGSDNRGSGRSY<br>NNDRRDNGGDGEFLVPRGSMVSKGEELFTGVVPILVELDGDVNGHKFSVS<br>GEGEGDATYGKLTLKFICTTGKLPVPWPTLVTTLTYGVQCFSRYPDHMKQHD<br>FFKSAMPEGYVQERTIFFKDDGNYKTRAEVKFEGDTLVNRIELKGIDFKEDG<br>NILGHKLEYNYNSHNVYIMADKQKNGIKVNFKIRHNIEDGSVQLADHYQQ<br>NTPIGDGPVLLPDNHYLSTQSALSKDPNEKRDHMVLLEFVTAAGITLGMDE<br>LYKEFLVPRGSMESNQSNNGGSGNAALNRGGRYVPPHLRGGDGGAAAAA<br>SAGGDDRRGGAGGGGYRRGGGNSGGGGGGGYDRGYNDNRDDRDNRG<br>GSGGYGRDRNYEDRGYNGGGGGGGNRGYNNNRGGGGGGYNRQDRGD<br>GGSSNFSRGGYNNRDEGSDNRGSGRSYNNDRRDNGGDGLEHHHHHH |
| MBP-RGG-GFP-RGG (uncleavable) | MMKIEEGKLVIWINGDKGYNGLAEVGKKFEKDTGIKVTVEHPDKLEEKFPQ<br>VAATGDGPDIIFWAHDRFGGYAQSGLLAEITPDKAFQDKLYPFTWDAVRYN<br>GKLIAYPIAVEALSLIYNKDLLPNPPKTWEEIPALDKELKAKGKSALMFNLQEP<br>YFTWPLIAADGGYAFKYENGKYDIKDVGVDNAGAKAGLTFLVDLIKNKHMN<br>ADTDYSIAEAAFNKGETAMTINGPWAWSNIDTSKVNYGVTVLPTFKGQPSK<br>PFVGVLSAGINAASPNKELAKEFLENYLLTDEGLEAVNKDKPLGAVALKSYEE<br>ELVKDPRIAATMENAQKGEIMPNIPQMSAFWYAVRTAVINAASGRQTVDE<br>ALKDAQHMESNQSNNGGSGNAALNRGGRYVPPHLRGGDGGAAAAASA<br>GGDDRRGGAGGGGYRRGGGNSGGGGGGGYDRGYNDNRDDRDNRGGS<br>GGYGRDRNYEDRGYNGGGGGGGNRGYNNNRGGGGGGYNRQDRGDGG<br>SSNFSRGGYNNRDEGSDNRGSGRSYNNDRRDNGGDGEFLVPRGSMVSKG<br>EELFTGVVPILVELDGDVNGHKFSVSGEGEGDATYGKLTLKFICTTGKLPVPW<br>PTLVTTLTYGVQCFSRYPDHMKQHDFFKSAMPEGYVQERTIFFKDDGNYKT<br>RAEVKFEGDTLVNRIELKGIDFKEDGNILGHKLEYNYNSHNVYIMADKQKNG<br>IKVNFKIRHNIEDGSVQLADHYQQNTPIGDGPVLLPDNHYLSTQSALSKDPN<br>EKRDHMVLLEFVTAAGITLGMDELYKEFLVPRGSMESNQSNNGGSGNAAL<br>NRGGRYVPPHLRGGDGGAAAAASAGGDDRRGGAGGGGYRRGGGNSGG<br>GGGGGYDRGYNDNRDDRDNRGGSGGYGRDRNYEDRGYNGGGGGGGN<br>RGYNNNRGGGGGGYNRQDRGDGGSSNFSRGGYNNRDEGSDNRGSGRSY<br>NNDRRDNGGDGLEHHHHHH |
| MBP-FUSLC-mCherry-RGG | MKIEEGKLVIWINGDKGYNGLAEVGKKFEKDTGIKVTVEHPDKLEEKFPQVA<br>ATGDGPDIIFWAHDRFGGYAQSGLLAEITPDKAFQDKLYPFTWDAVRYNGK |

| | |
|---|---|
| | LIAYPIAVEALSLIYNKDLLPNPPKTWEEIPALDKELKAKGKSALMFNLQEPYF<br>TWPLIAADGGYAFKYENGKYDIKDVGVDNAGAKAGLTFLVDLIKNKHMNA<br>DTDYSIAEAAFNKGETAMTINGPWAWSNIDTSKVNYGVTVLPTFKGQPSKP<br>FVGVLSAGINAASPNKELAKEFLENYLLTDEGLEAVNKDKPLGAVALKSYEEE<br>LVKDPRIAATMENAQKGEIMPNIPQMSAFWYAVRTAVINAASGRQTVDEA<br>LKDAQTGGGGSGGGGSLEVLFQGPGHMASNDYTQQATQSYGAYPTQPG<br>QGYSQQSSQPYGQQSYSGYSQSTDTSGYGQSSYSSYGQSQNTGYGTQSTP<br>QGYGSTGGYGSSQSSQSSYGQQSSYPGYGQQPAPSSTSGSYGSSSQSSSYG<br>QPQSGSYSQQPSYGGQQQSYGQQQSYNPPQGYGQQNQYNSGSMVSKG<br>EEDNMAIIKEFMRFKVHMEGSVNGHEFEIEGEGEGRPYEGTQTAKLKVTKG<br>GPLPFAWDILSPQFMYGSKAYVKHPADIPDYLKLSFPEGFKWERVMNFEDG<br>GVVTVTQDSSLQDGEFIYKVKLRGTNFPSDGPVMQKKTMGWEASSERMY<br>PEDGALKGEIKQRLKLKDGGHYDAEVKTTYKAKKPVQLPGAYNVNIKLDITS<br>HNEDYTIVEQYERAEGRHSTGGMDELYKGGGSENLYFQGEFGKLMESNQS<br>NNGGSGNAALNRGGRYVPPHLRGGDGGAAAAASAGGDDRRGGAGGGG<br>YRRGGGNSGGGGGGGYDRGYNDNRDDRDNRGGSGGYGRDRNYEDRGY<br>NGGGGGGGNRGYNNNRGGGGGGYNRQDRGDGGSSNFSRGGYNNRDE<br>GSDNRGSGRSYNNDRRDNGGDGAAALEHHHHHH |
| MBP-FUSLC-GST-mCherry-RGG | MKIEEGKLVIWINGDKGYNGLAEVGKKFEKDTGIKVTVEHPDKLEEKFPQVA<br>ATGDGPDIIFWAHDRFGGYAQSGLLAEITPDKAFQDKLYPFTWDAVRYNGK<br>LIAYPIAVEALSLIYNKDLLPNPPKTWEEIPALDKELKAKGKSALMFNLQEPYF<br>TWPLIAADGGYAFKYENGKYDIKDVGVDNAGAKAGLTFLVDLIKNKHMNA<br>DTDYSIAEAAFNKGETAMTINGPWAWSNIDTSKVNYGVTVLPTFKGQPSKP<br>FVGVLSAGINAASPNKELAKEFLENYLLTDEGLEAVNKDKPLGAVALKSYEEE<br>LVKDPRIAATMENAQKGEIMPNIPQMSAFWYAVRTAVINAASGRQTVDEA<br>LKDAQTGGGGSGGGGSLEVLFQGPGHMASNDYTQQATQSYGAYPTQPG<br>QGYSQQSSQPYGQQSYSGYSQSTDTSGYGQSSYSSYGQSQNTGYGTQSTP<br>QGYGSTGGYGSSQSSQSSYGQQSSYPGYGQQPAPSSTSGSYGSSSQSSSYG<br>QPQSGSYSQQPSYGGQQQSYGQQQSYNPPQGYGQQNQYNSGSMSPILG<br>YWKIKGLVQPTRLLLEYLEEKYEEHLYERDEGDKWRNKKFELGLEFPNLPYYI<br>DGDVKLTQSMAIIRYIADKHNMLGGCPKERAEISMLEGAVLDIRYGVSRIAYS<br>KDFETLKVDFLSKLPEMLKMFEDRLCHKTYLNGDHVTHPDFMLYDALDVVL<br>YMDPMCLDAFPKLVCFKKRIEAIPQIDKYLKSSKYIAWPLQGWQATFGGGD<br>HPPGSGMVSKGEEDNMAIIKEFMRFKVHMEGSVNGHEFEIEGEGEGRPYE<br>GTQTAKLKVTKGGPLPFAWDILSPQFMYGSKAYVKHPADIPDYLKLSFPEGF<br>KWERVMNFEDGGVVTVTQDSSLQDGEFIYKVKLRGTNFPSDGPVMQKKT<br>MGWEASSERMYPEDGALKGEIKQRLKLKDGGHYDAEVKTTYKAKKPVQLP<br>GAYNVNIKLDITSHNEDYTIVEQYERAEGRHSTGGMDELYKGGGSENLYFQG<br>EFGKLMESNQSNNGGSGNAALNRGGRYVPPHLRGGDGGAAAAASAGGD<br>DRRGGAGGGGYRRGGGNSGGGGGGGYDRGYNDNRDDRDNRGGSGGY<br>GRDRNYEDRGYNGGGGGGGNRGYNNNRGGGGGGYNRQDRGDGGSSN<br>FSRGGYNNRDEGSDNRGSGRSYNNDRRDNGGDGAAALEHHHHHH |

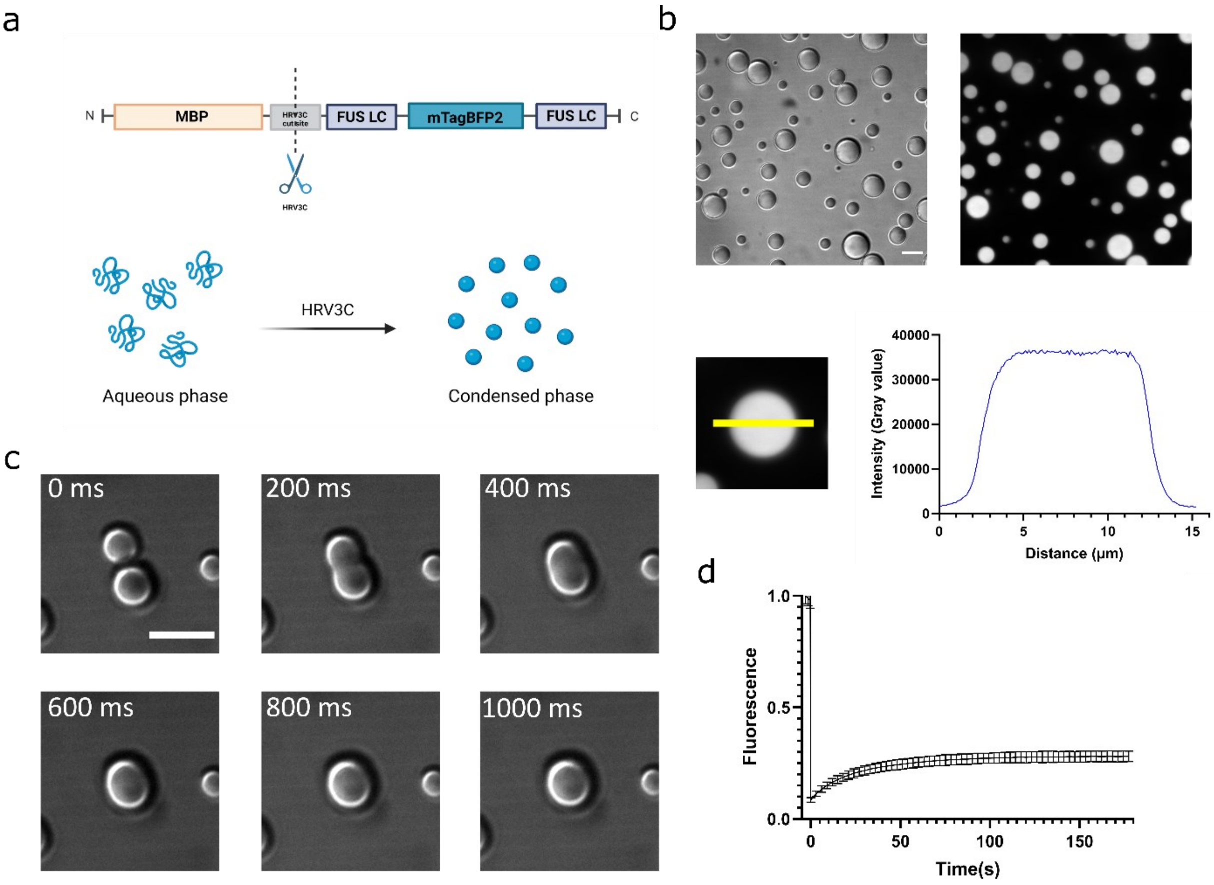


**Fig. S1. Designing FUSLC condensates with controlled formation. 1a.** Illustration of FUS LC condensates design. When attached to MBP, the protein remains in the aqueous phase. Upon cleaveage of MBP by HRV-3C protease, FUSLC-mTagBFP2-FUSLC phase separates to form a condensed phase. **1b.** FUSLC-mTagBFP2-FUSLC forms micron sized spherical droplets in vitro. Upper left, DIC image; upper right, fluorescence image. Lower panel, fluorescence intensity profile of FUSLC-mTagBFP2-FUSLC droplet. MBP-FUSLC-mTagBFP2-FUSLC protein was incubated with HRV-3C protease at 42 ℃ before being added into the imaging chamber for confocal imaging. 7.5 μM protein final concentration in a buffer of 20 mM HEPES, 150 mM NaCl, pH 7.4. Scale bar: 10 μm. **1c.** Fusion of FUSLC-mTagBFP2-FUSLC droplets. DIC micrographs were taken with 50 ms exposure and 150 ms interval in a timelapse series. Scale bar: 10 μm. **1d.** FRAP curve for FUSLC-mTagBFP2-FUSLC droplets. MBP-FUSLC-mTagBFP2-FUSLC was incubated with HRV-3C protease at 42 ℃ before being added into the imaging chamber. 7.5 μM protein final concentration in 20 mM HEPES, 150 mM NaCl, pH 7.4. The droplets were bleached by a 405 nm laser in circular 1 μm diameter ROIs and fluorescence recovery was tracked over 180 s. n = 10. Scale bar: 10 μm.

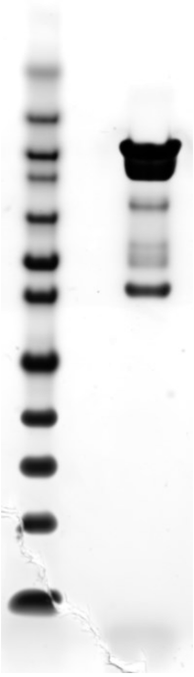

**Fig. S2** Protein gel for MBP-FUSLC-mTagBFP2-FUSLC. Expected molecular weight: 104.9 kDa. Protein standard bands from top to bottom: 260, 160, 110, 80, 60, 50, 40, 30, 20, 15, 10, and 3.5 kDa.

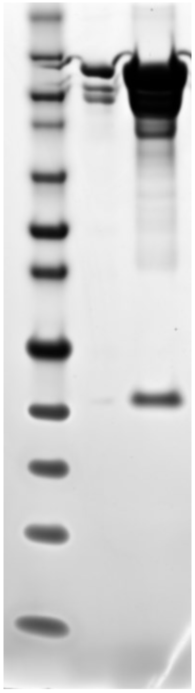

**Fig. S3** Protein gel for MBP-FUS $_{FL}$-mCherry. Expected molecular weight: 123.4 kDa. Protein standard bands from top to bottom: 260, 160, 110, 80, 60, 50, 40, 30, 20, 15, 10, and 3.5 kDa.

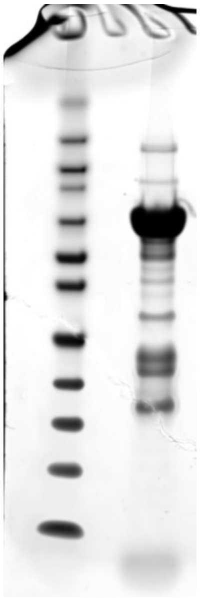

**Fig. S4** Protein gel for RGG-GFP-RGG. Expected molecular weight: 63.3 kDa. Protein standard bands from top to bottom: 260, 160, 110, 80, 60, 50, 40, 30, 20, 15, 10, and 3.5 kDa.

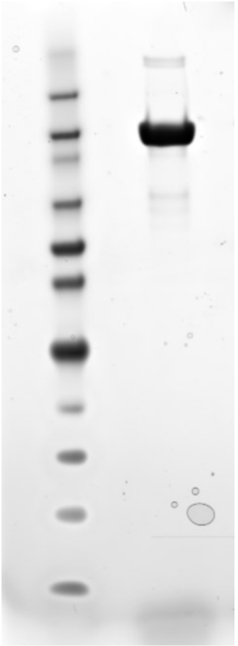

**Fig. S5** Protein gel for MBP-RGG-GFP-RGG (HRV-3C site). Expected molecular weight: 105.7 kDa. Protein standard bands from top to bottom: 260, 160, 110, 80, 60, 50, 40, 30, 20, 15, 10, and 3.5 kDa.

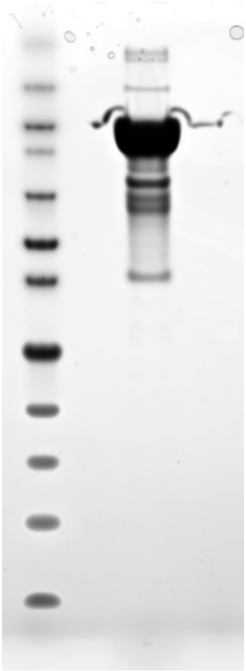

**Fig. S6** Protein gel for MBP-RGG-GFP-RGG (TEV site). Expected molecular weight: 106.2 kDa. Protein standard bands from top to bottom: 260, 160, 110, 80, 60, 50, 40, 30, 20, 15, 10, and 3.5 kDa.

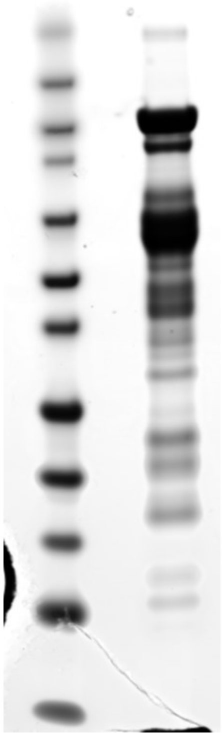

**Fig. S7** Protein gel for MBP- RGG-GFP-A$\beta_{42}$-RGG. Expected molecular weight: 110.5 kDa. Protein standard bands from top to bottom: 260, 160, 110, 80, 60, 50, 40, 30, 20, 15, 10, and 3.5 kDa.

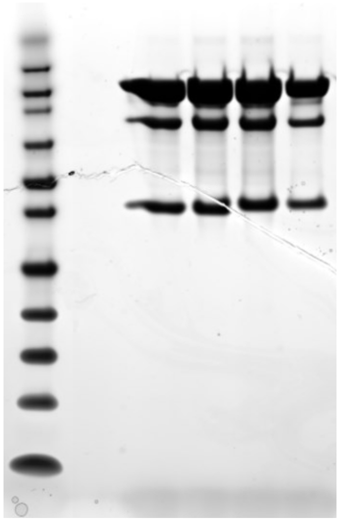

**Fig. S8** Protein gel for MBP-FUSLC-mCherry-RGG (4 aliquots). Expected molecular weight: 105.8 kDa. Protein standard bands from top to bottom: 260, 160, 110, 80, 60, 50, 40, 30, 20, 15, 10, and 3.5 kDa.

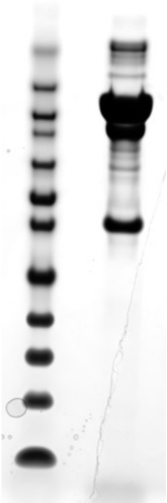

**Fig. S9** Protein gel for MBP-FUSLC-GST-mCherry-RGG. Expected molecular weight: 131.3 kDa. Protein standard bands from top to bottom: 260, 160, 110, 80, 60, 50, 40, 30, 20, 15, 10, and 3.5 kDa.

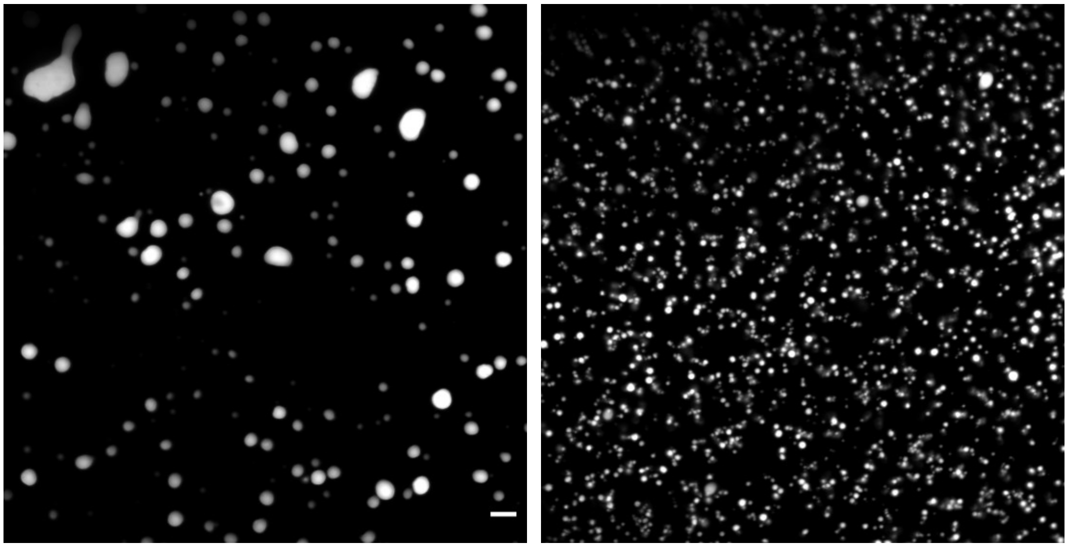

**Fig. S10** FUSLC-mTagBFP2-FUSLC droplets after adding high concentration of salt solution. 4 M Final NaCl concentration. Left, immediately after adding salt solution, droplets were deformed due to shear force, but not dissolved. Right, 1 day after, smaller gel like droplets were seen. 10 µm.

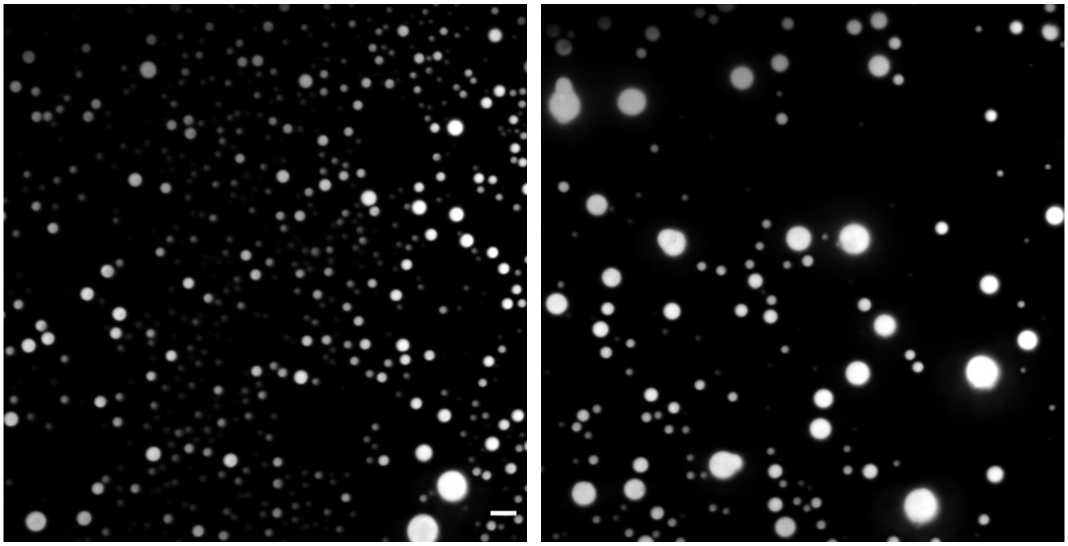

**Fig. S11** FUSLC-mTagBFP2-FUSLC droplets at longer incubation at room temperature. Left, droplets at 48 hr. Right, droplets at 120 hr. Scale bar: 10 µm.

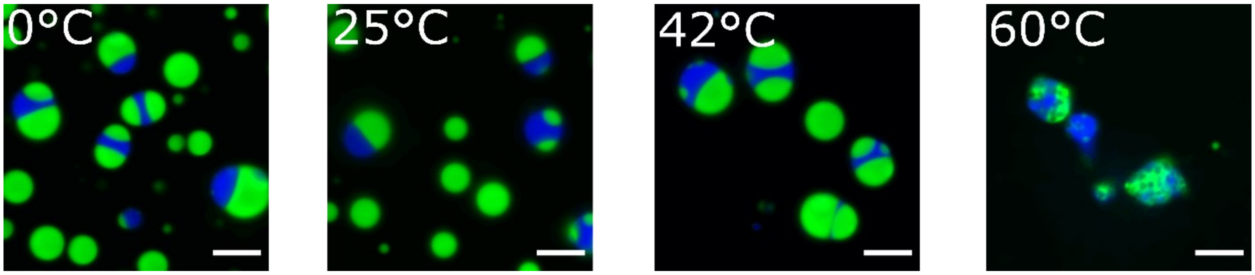


**Fig. S12** Effect of MBP-RGG-GFP-RGG incubation temperature on FUS LC / RGG mixing structure. MBP-FUSLC-mTagBFP2-FUSLC and HRV-3C were incubated at 42 °C before being diluted into imaging well. MBP-RGG-GFP-RGG and HRV-3C were incubated with HRV-3C at different temperatures before being added into the wells. Incubation at 60 °C appeared to have denatured the RGG-GFP-RGG protein and negatively affected the RGG condensates. Overall, the two phases remain immiscible and partially wetted at these temperatures. Images were acquired at room temperature. Scale bar: 10 µm. Incubation of FUSLC-mTagBFP2-FUSLC droplets at lower temperature (0 °C) promoted FUS LC droplets to form gel-like blobs.

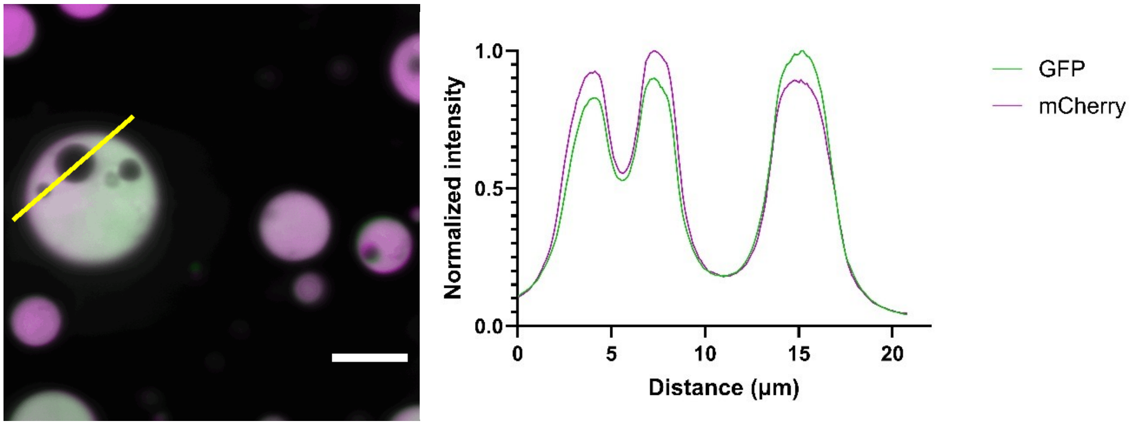


**Fig. S13** Kinetically trapped droplets formed, after first forming RGG-GFP-RGG condensates in wells, then adding FUS $_{FL}$-mCherry. 5.5 µM of each protein final concentration. Scale bar: 10 µm.

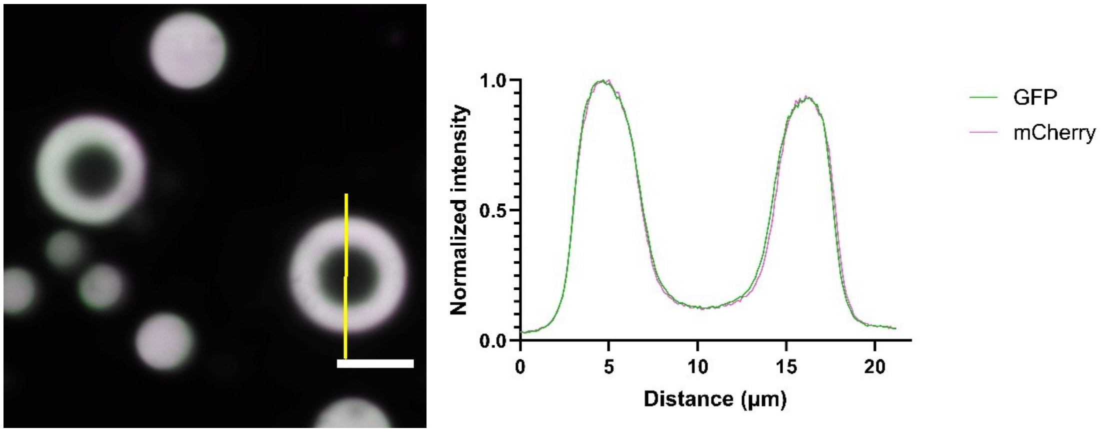


**Fig. S14** Kinetically trapped droplets formed, after pre-mixing RGG-GFP-RGG with MBP-FUS $_{FL}$-mCherry and HRV-3C at 42 °C before being diluted into imaging well. 5.5 μM of each protein final concentration. Scale bar: 10 μm.

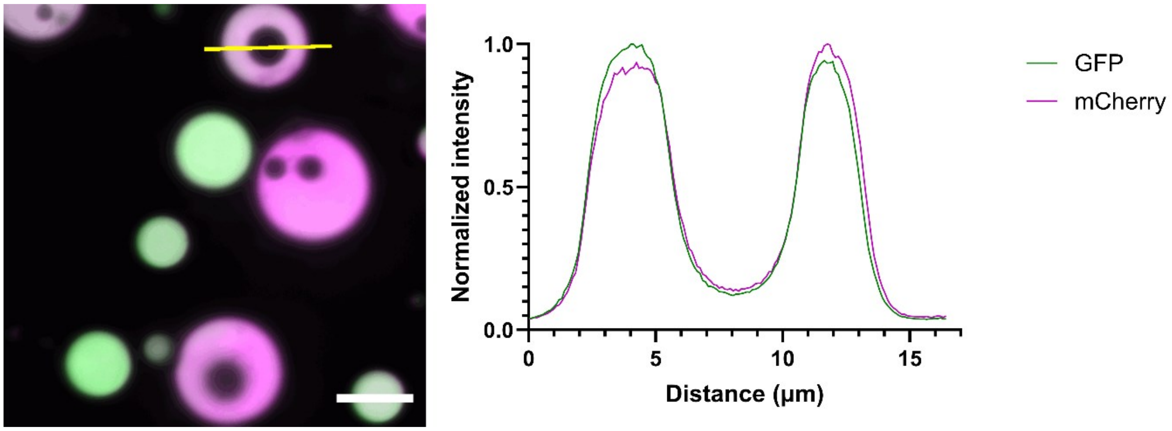


**Fig. S15.** Kinetically trapped droplets formed, after first forming FUS $_{FL}$-mCherry condensates in wells, then adding RGG-GFP-RGG. 5.5 μM of each protein final concentration. Scale bar: 10 μm.

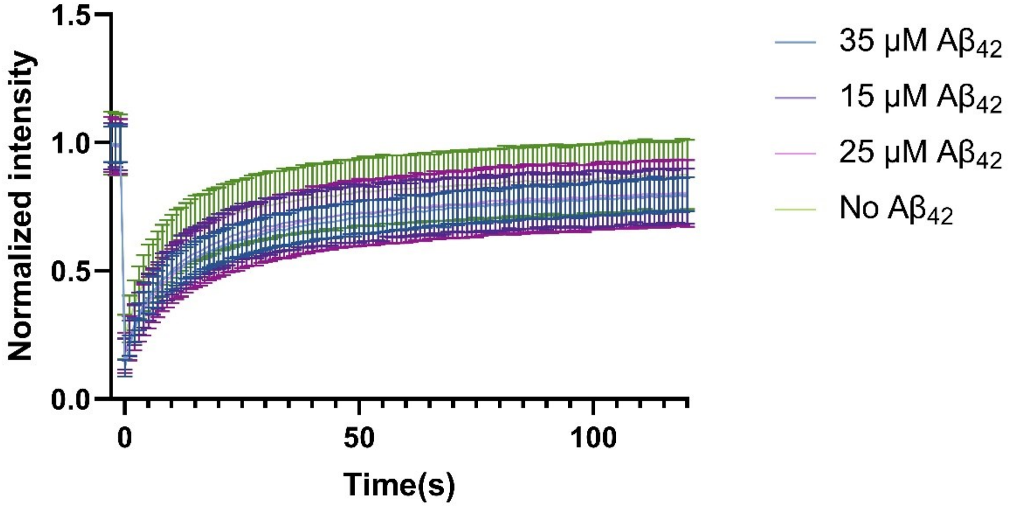


**Fig. S16.** FRAP curve on RGG-GFP-RGG condensates with varying concentration of free Aβ$_{42}$ peptide added to the well. Incubated for 1 day. Regions of 1 μm diameter bleached by 405 nm laser. Recovery measured for 120 s at 1 s interval.

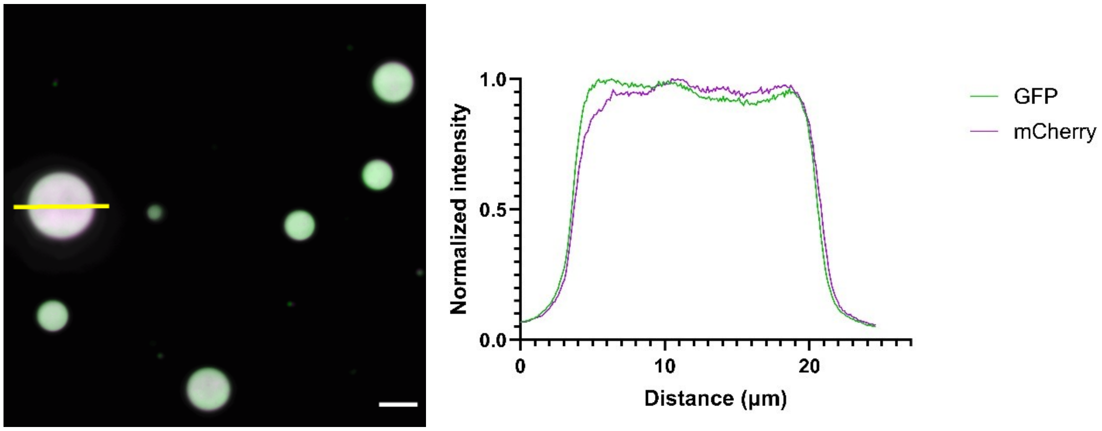


**Fig. S17.** Premixing FUS$_{FL}$ with RGG-GFP-Aβ$_{42}$-RGG overcame the miscibility barrier caused by viscosity. MBP-RGG-GFP-Aβ$_{42}$-RGG and MBP-FUS$_{FL}$-mCherry were co-incubated with HRV-3C protease before diluting into the imaging well. 9 μM RGG-GFP-Aβ$_{42}$-RGG and 6.6 μM FUS$_{FL}$-mCherry final concentration. Scale bar: 10 μm.

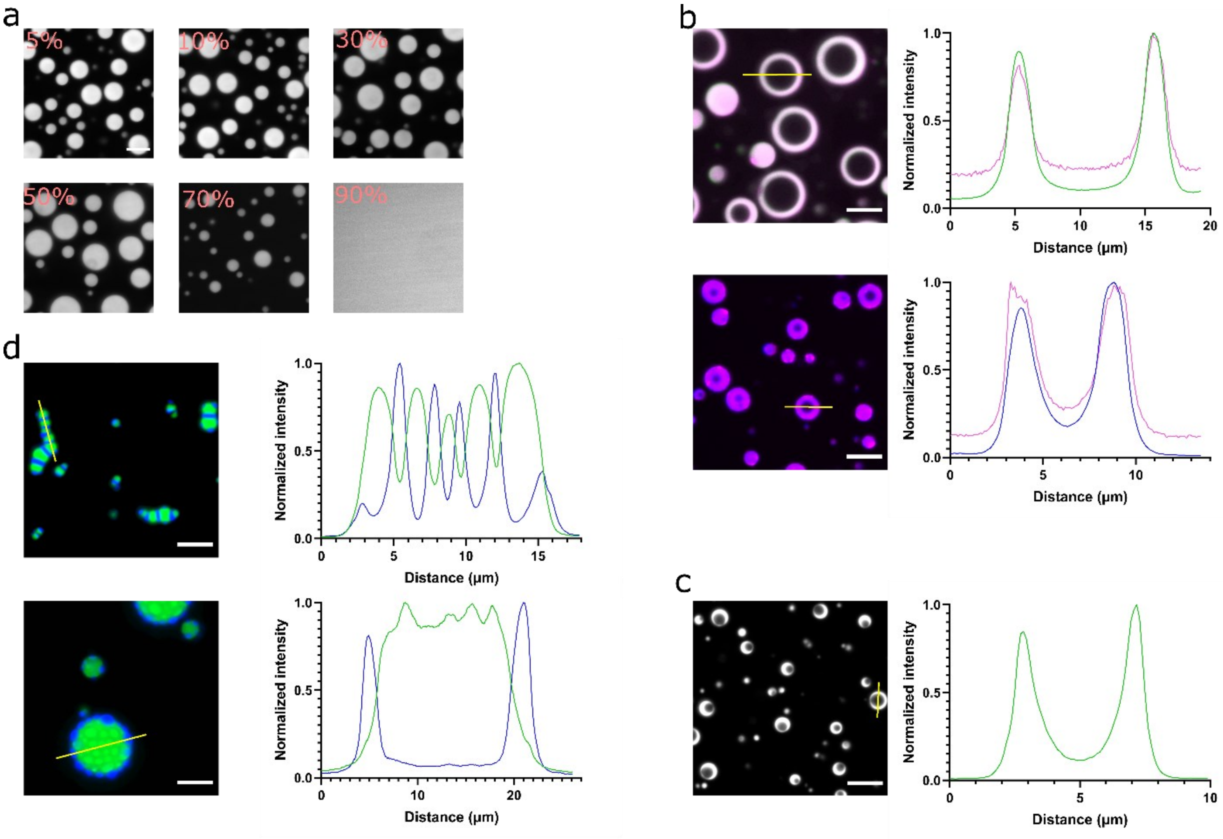


**Fig. S18.** Control experiments on formation of kinetically trapped droplets. **a.** Kinetically trapped double-emulsion droplets are not a result of incomplete protease digestion. Co-incubation of cleavable and uncleavable versions of MBP-RGG-GFP-RGG at different ratios with HRV-3C at 42 ℃ for 20 min before being diluted into the imaging well. The uncleavable version of the protein does not contain the HRV-3C cut site. Images show an uncleavable composition of 5%, 10%, 30%, 50%, 70%, and 90%, respectively. 18 μM total protein concentration in 20 mM HEPES, 150 mM NaCl, PH 7.4. Scale bar: 10 μm. **b.** HRV-3C protease does not stabilize the interfaces of double-emulsion droplets. HRV-3C labelled by Alexa Fluor 594 C5 maleimide (magenta) colocalizes with RGG-GFP-RGG (green, top panel) and FUSLC-mTagBFP2-FUSLC (blue, lower panel) phases. Scale bar: 10 μm. **c.** Formation of double-emulsion droplets is not specific to HRV-3C protease. Replacing HRV-3C with TEV also yielded kinetically trapped droplets. 10 μl 36 μM MBP-RGG-GFP-RGG (with TEV cut site) and 0.1 μl 0.2 mg/ml proTEV protease was co-incubated at 42 ℃ for 30 s then 5 μl of the mixture was diluted 6-fold in 20 mM HEPES buffer, PH 7.4 for imaging. Scale bar: 10 μm. **d.** Co-incubation of RGG-GFP-RGG (no MBP) with MBP-FUSLC-mTagBFP2-FUSLC and HRV-3C at 42 ℃ before being diluted into imaging well. Top: 7.2 μM RGG-GFP-RGG and 3.6 μM FUSLC-mTagBFP2-FUSLC. Lower: 9 μM RGG-GFP-RGG and 3 μM FUSLC-mTagBFP2-FUSLC. 20 mM HEPES, 150 mM NaCl, PH 7.4. Scale bar: 10 μm.

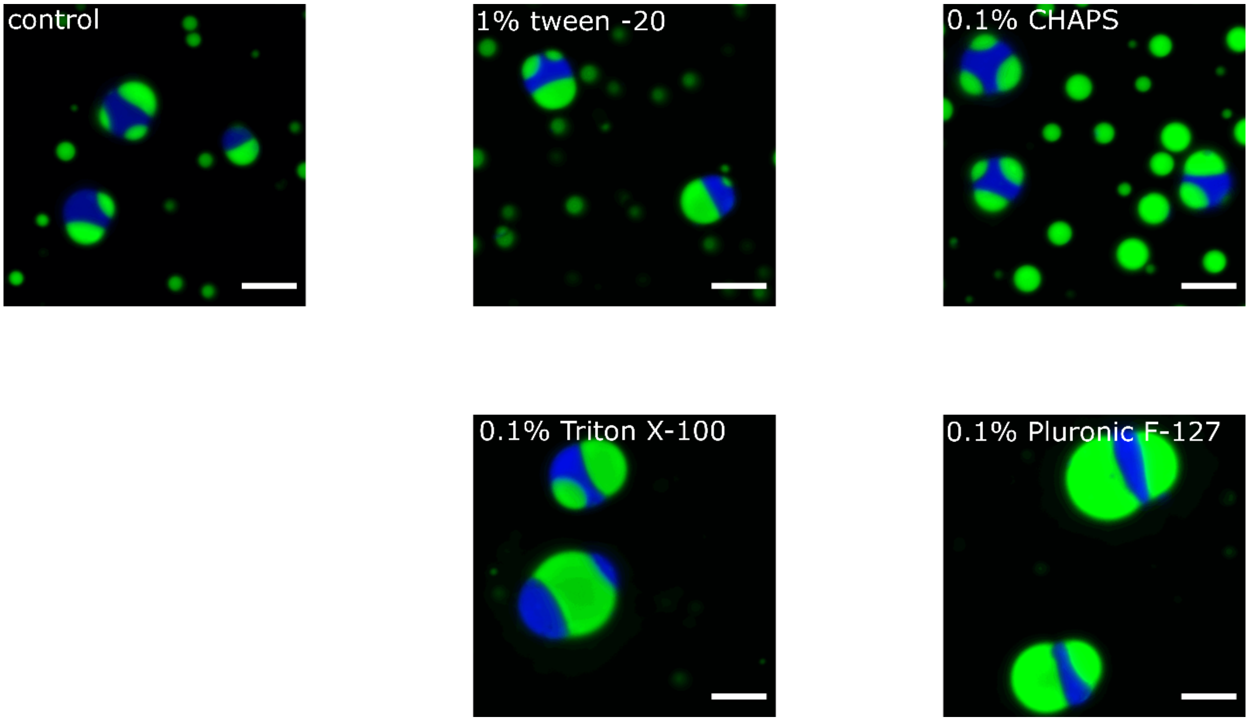


**Fig. S19.** Testing effect of conventional surfactants on FUS LC / RGG condensates mixture structure. Some of the common surfactants do not affect the partially wetted structure of mixed FUS LC / RGG condensates. MBP-FUSLC-mTagBFP2-FUSLC and HRV-3C were incubated at 42 °C before being diluted into imaging well. MBP-RGG-GFP-RGG and HRV-3C were then incubated with HRV-3C at 42°C before being added into the wells. Scale bars: 10 µm.

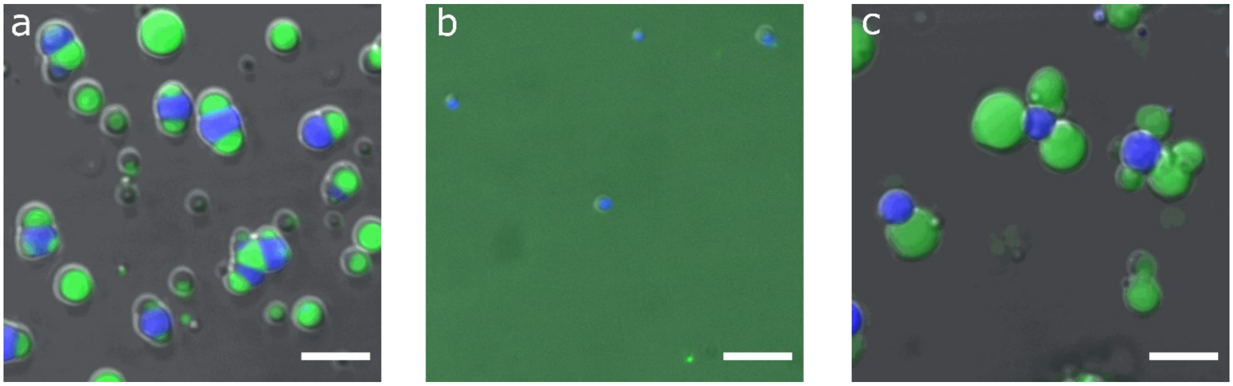


**Fig. S20.** Reentrant phase transition induced by PolyU RNA. **a**, FUS LC and RGG condensates before addition of RNA. **b**, addition of RNA dissolved condensates. Final RNA concentration of 800 nM. **c**, reentrant phase transition of FUS LC and RGG condensates. Images are merge of DIC (gray), 488 nm (green), and 405 nm (blue) channels. Scale bars: 10 µm.

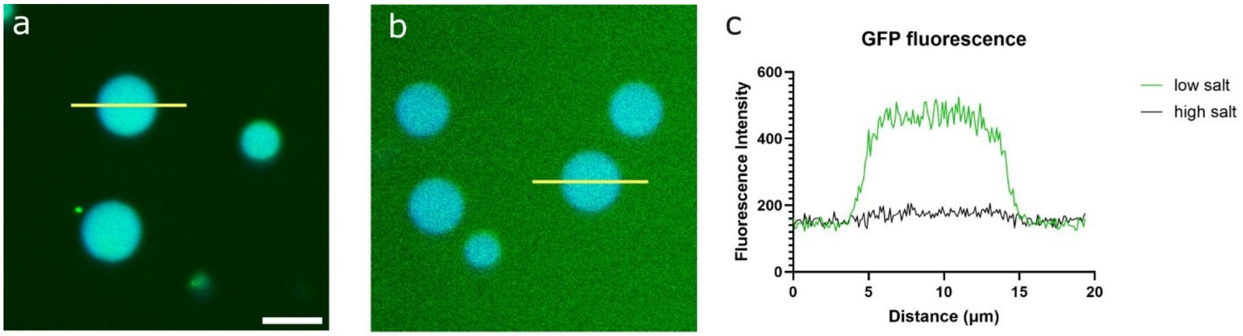


**Fig. S21.** RGG-GFP-RGG peptide at non-phase separating concentration partitions into the FUS LC phase. **a,** at physiological salt concentration, RGG-GFP-RGG at 300 nM concentration (below the saturation concentration) partitions into the FUS LC phase. **b,** raising salt concentration in the same imaging well by 500 mM favored RGG-GFP-RGG partitioning into the outer aqueous phase. **c,** fluorescence intensity profile at different salt concentrations. Scale bars: 10 µm.
